# Dynamic neuronal activation of a distributed cortico-basal ganglia-thalamus loop in learning a delayed sensorimotor task

**DOI:** 10.1101/568055

**Authors:** Xiaowei Gu, Chengyu T. Li

## Abstract

The cortico-basal ganglia-thalamus (CBT) loop is important for behavior. However, the activity and learning-related modulation within the loop in behavior remain unclear. To tackle this problem, we trained mice to perform a delayed sensorimotor-transformation task and recorded single-unit activity during learning simultaneously from four regions in a CBT loop: prelimbic area (PrL), posterior premotor cortex (pM2), dorsomedial caudate/putamen (dmCP), and mediodorsal thalamus (MD). Sensory and decision related information were encoded by the neurons within the loop, with weak interaction among neurons of different coding ability. The functional interaction among regions within the loop was dynamically routed in the loop during different behavioral phases and contributed to explain decision-related neuronal activity. The neurons of PrL and dmCP exhibited learning-related reorganization in neuronal activity and more persistent coding of sensory and decision-related information. Thus, both sensory- and decision-related information are processed in a functionally interacted CBT loop that is modulated by learning.

---

The cortex-basal ganglia-thalamus cortical (CBT) loop plays a central role to adaptive behavioral control *(1–8)*. Anatomically, cortex and thalamus send excitatory projections to caudate/striatum of basal ganglia, which recurrently and topographically project back to cortex through intermediate regions including basal ganglia and thalamus *(1, 8)*. Perturbation and recording studies demonstrated that the regions in the loop play critical roles in many processes, including motor control*(2–4, 6, 9, 10)*, reinforcement*(11–13)*, perceptual decision making*(14–16)*, inhibitory control*(5, 17)*, and working memory*(18–31)*. Furthermore, impaired functions of the loop has been implicated in many psychiatric diseases*(7)*. Therefore, it is important to understand how neurons within this CBT loop work together. However, the recurrent nature of the connectivity of this loop poses a notable challenge for understanding behavior-relevant coordination in its population activity.

We focused on a behavioral task requiring working memory (WM) and sensorimotor transformation (SMT). WM is a process of the brain to actively maintain and manipulate information for a brief delay period of several seconds to guide behavior *(32, 33)*. SMT is a process of transforming sensory inputs to a motor output based on behavioral context. It is well-known that prefrontal, parietal, and motor-related cortical areas, basal ganglion and thalamus are involved in WM *(19, 34)* and SMT *(35–38)*. However, recording from different regions were typically obtained from different subjects in different behavioral tasks, rendering difficulties in comparing the results. A simple hypothesis explaining the relationship of WM and SMT related activity is that single neurons can code for both the sample and test odors, in a manner related with pairing relationship (as in delayed match to sample task*(41)*. However, it is unclear whether this simple hypothesis can explain the relationship between WM- and SMT-related activities in the CBT loop. Furthermore, functional interaction among brain regions may undergo distinct dynamics in different phases of learning. But only a few studies recorded neuronal activity throughout learning period of WM tasks *(27, 39, 40)*.

We hereby trained head-fixed mice to perform an olfactory delayed paired association task. In the ODPA task, mice need to maintain sensory WM during a delay period and then decide whether to lick based on specific pairing of sample and test odors. Therefore, the maintained WM needs to be integrated and compared with the upcoming sensory information, then transformed into a lick or no-lick decision. By simultaneously recording single-neuron activity from four regions in a CBT loop, we found distributed representation of both sensory- and decision-related information within a CBT loop and its dynamics in cross-region interaction through learning.

## Behavioral paradigm

We trained head-fixed mice to perform a behavioral task based on an automatic training system (**Fig. 1A**, Sfig. 1)*(42)*. Specifically mice were trained with an olfactory delayed pair-association (DPA) task, as in *(43)*. Mice need to maintain the sensory information of a sample odor (S1 or S2) during the delay period (5 sec in duration), after which a test odor (T1 or T2) was delivered (**Fig. 1B**). Licking within a response window in the paired trials (S1-T1 or S2-T2) leads to water reward (scored as hit if mice licked, as miss otherwise), but not in the unpaired trials (S1-T2 or S2-T1, scored as false alarm if mice licked, as correct rejection otherwise). Mice learnt the task well within several days (**Fig. 1C**). The learning was manifested as the increased correct rejection rate in the unpaired trials (SFig. 2A-B). Throughout the learning process, mice maintained a high lick probability in the paired trials (SFig. 2C-D). Mice did not lick during either sample-odor delivery or delay period (SFig. 2E).

**Figure 1:**
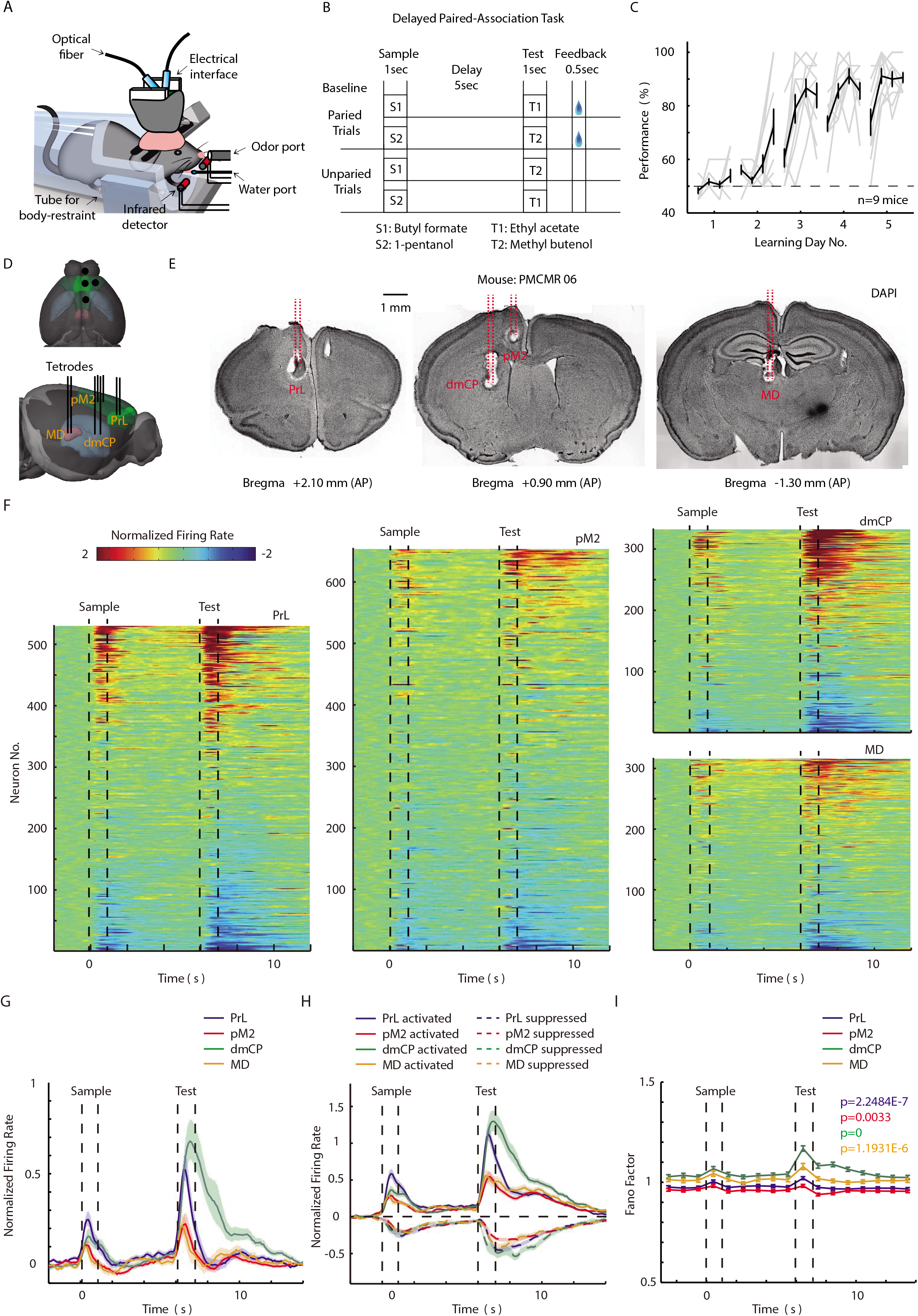
Behavior paradigm and distributed modulation in the neuronal activity within the CBT loop of the olfactory DPA task. (**A**) Schematic of training facility for mice performing olfactory tasks. (**B**) Protocol of mice performing olfactory delayed pair-association (DPA) task. (**C**) Overall performance of mice in learning olfactory DPA task from day1 to day5. (**D**) A diagram of the recording sites and their spatial corresponding relation. (**E**) The brain slices of thermo lesion sites at the tip of electrodes indicate the recording sites at the four brain regions. (**F**) Heat map of normalized firing rate for all neurons recorded in four regions. Each row represents the baseline-normalized firing rate of one neuron (color bar indicated for the Z-scores normalization calculated from the cross-trial mean and standard deviation during the baseline period). The y-axis labels represent brain regions and the number of neurons recorded in the corresponding region. The two dotted lines represent the onset and offset of odor delivery. Time zero represents the onset of sample-odor delivery. Normalized firing rate is defined as raw firing rate subtracting the mean of baseline firing across all trials and dividing the standard deviation of baseline firing across trials. It is noted that this normalized firing rate represent firing modulation of neurons during task events. (**G**) Averaged normalized firing rate of neurons from PrL, pM2, dmCP and MD. (**H**) Averaged normalized firing rate of positively and negatively modulated neurons from PrL, pM2, dmCP and MD. (**I**) Famo factors of normalized firing rate in each 100ms bin for neurons from PrL, pM2, dmCP and MD.

### Distributed modulation in neuronal activity within the CBT loop by the task stimuli

We custom-made tetrode with twisted wire to perform single-unit recording simultaneously from four regions of a CBT loop: PrL, pM2, dmCP and MD (**Fig. 1D**, SFig. 3). The recording locations were verified by electrical lesion after recording (**Fig. 1E**).

In our multi-region recordings, typically about ten neurons were recorded simultaneously in each region on each day of recording (see materials and methods, SFig. 4). Overall 1837 neurons were recorded. To visualize the activity modulation of each region in the task, we plotted the Z-score normalized firing rate of each neuron (**Fig. 1F**). The neuronal activity of all four regions was modulated during the sample-delivery, delay, and response periods (**Fig. 1F**, SFig. 5). Both activation and suppression in neuronal activity were observed following the sample- and test-odor delivery (cells in red and blue in **Fig. 1F**). On average there was a transient increase in the population firing rate during the odor-delivery period (**Fig. 1G**), then a small ramping-up modulation during delay period, as in *(44)*. We then separated the neurons into the activated and suppressed groups. For the activated group, the neurons of PrL and dmCP are more strongly modulated by the sample and test odors (**Fig. 1H**). For the negatively modulated group, the difference of modulation among the four regions was not statistically significant (**Fig. 1H**). Variance in neuronal firing was increased during the odor-delivery period, as revealed by the Fano-factor analysis (**Fig. 1I**).

### Neurons of the CBT loop maintained task-related sensory information

A hallmark of WM-related activity is the ability of coding the maintained information during the delay period *(19, 24, 26, 27, 30, 34, 43–46)*. The behavioral design of the DPA task is sensory oriented, because motor-related decision cannot be made during delay period. Therefore, one can predict that an action-oriented CBT loop may not necessarily maintain the sensory information during the delay period. Instead, we observed many neurons in all four regions of this loop maintaining the sensory information (**Fig. 2A-C**). An example neuron with different firing rates following different sample odors was plotted in Fig. 2A. We defined the sensory selectivity of a neuron as the difference between the average firing rate following different sample odors, divided by the sum. We then plotted the sample-odor selectivity from all four regions in the heat maps to visualize the overall distribution of the sample selectivity (**Fig. 2B**). For each region, the average sample selectivity for the neurons preferring S1 or S2 was balanced (**Fig. 2C**). Both the selectivity for individual neurons (heat-map, **Fig. 2B**) and averaged population (**Fig. 2C**) demonstrated that the four regions exhibited different levels of selectivity for sample odor stimulus. The neurons in PrL and dmCP exhibited stronger sample selectivity in the early delay period, whereas the neurons in dmCP exhibited higher selectivity in the late delay period (SFig. 6). To verify the selectivity results, we performed the decoding analysis based on the classifier of maximum correlation coefficient (MCC)*(47)*. The sample information can be successfully decoded during the early delay period in all four regions and the late delay period in PrL, pM2, and dmCP (**Fig. 2D**).

**Figure 2:**
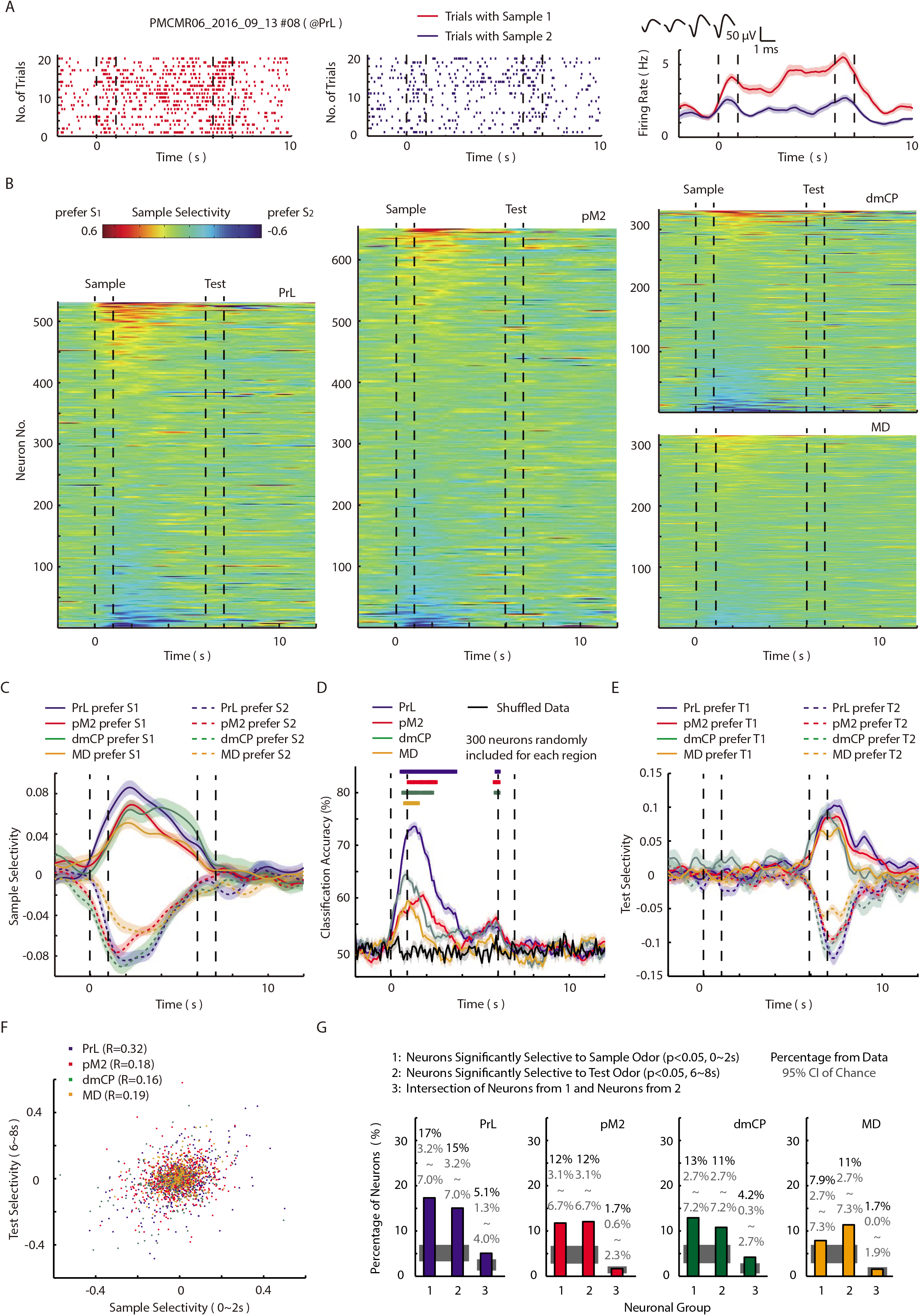
Neurons of the CBT loop maintained task-related sensory information. (**A**) Spiking raster plot (left and mid) and peristimulus time histogram (PSTH, right) showing a neuron in PrL with different activity during the sample-delivery and delay periods in trials following sample 1 (in red) and sample 2 (in blue). (**B**) Heat map of sample selectivity for all neurons recorded in the four regions. Sample selectivity for each neuron at certain time bin is defined as the difference of average firing rate at this bin from trial with S1 and S2, divided by the summation of them as normalization (Sample selectivity = (Activity^S1^ - Activity^S2^) / (Activity^S1^ + Activity^S2^)). (**C**) Averaged selectivity for sample odors of neurons preferring S1 or S2 from PrL, pM2, dmCP and MD. (**D**) Decoding accuracy of sample odors based on the classifier of maximum correlation coefficient for neurons in PrL, pM2, dmCP and MD. The colored squares on top indicate the significant difference from shuffled data for corresponding curves in the 100ms bin. (**E**) Averaged selectivity for test odors of neurons preferring T1 or T2 from PrL, pM2, dmCP and MD. (**F**) Correlation between selectivity for sample odors during 0~2s and selectivity for test odors during 6~8s for neurons in PrL, pM2, dmCP and MD. (**G**) Percentage of neurons selective to sample odors, test odors or both in PrL, pM2, dmCP and MD.

We then calculated the selectivity for the test odor. The neurons in all four regions significantly coded for the test odor (test-odor selectivity in **Fig. 2E**; decoding in SFig. 6C). To test the relationship between the ability of the CBT loop in coding the sample and test odors, we plotted the test-odor selectivity against the sample-odor selectivity. We observed significant but small correlation between the sample- and test-odor selectivity in all four regions (**Fig. 2F**). We also observed higher-than-chance level overlap in the neurons with significant sample- and test-odor selectivity in PrL and dmCP (**Fig. 2G**). Therefore, the coding of odor information in the CBT loop is weakly correlated for sample and test odors.

### Neurons of the CBT loop represented the pairing-related information

To accomplish this olfactory memory task, mice need to integrate the information of sample and test odors across the delay period to make a licking decision, according to paired/non-paired relationship. The presence of both sample- and test-odor information in the CBT loop (**Fig. 2**) suggested the possibility that the computation within the loop might be calculating the pairing relationship. We therefore analyzed the coding ability of the pairing relationship and the process of SMT in the CBT loop.

We firstly analyzed the coding ability for the pairing relationship in the individual neurons of the four regions. An example neuron was plotted showing different firing rates following the paired and unpaired odor stimuli (**Fig. 3A**). The selectivity for pairing relationship was defined as the difference between the average firing rate in the trials with the paired and unpaired odors, divided by the sum. The neurons of all four regions exhibited significant pairing selectivity (**Fig. 3B-C**). There was no significant pairing selectivity before the test-odor delivery (**Fig. 3C**), consistent with the lack of pairing information during this period. During the test-odor delivery and response periods, the neurons in PrL and dmCP exhibited stronger decoding power for pairing relationship (**Fig. 3D**).

**Figure 3:**
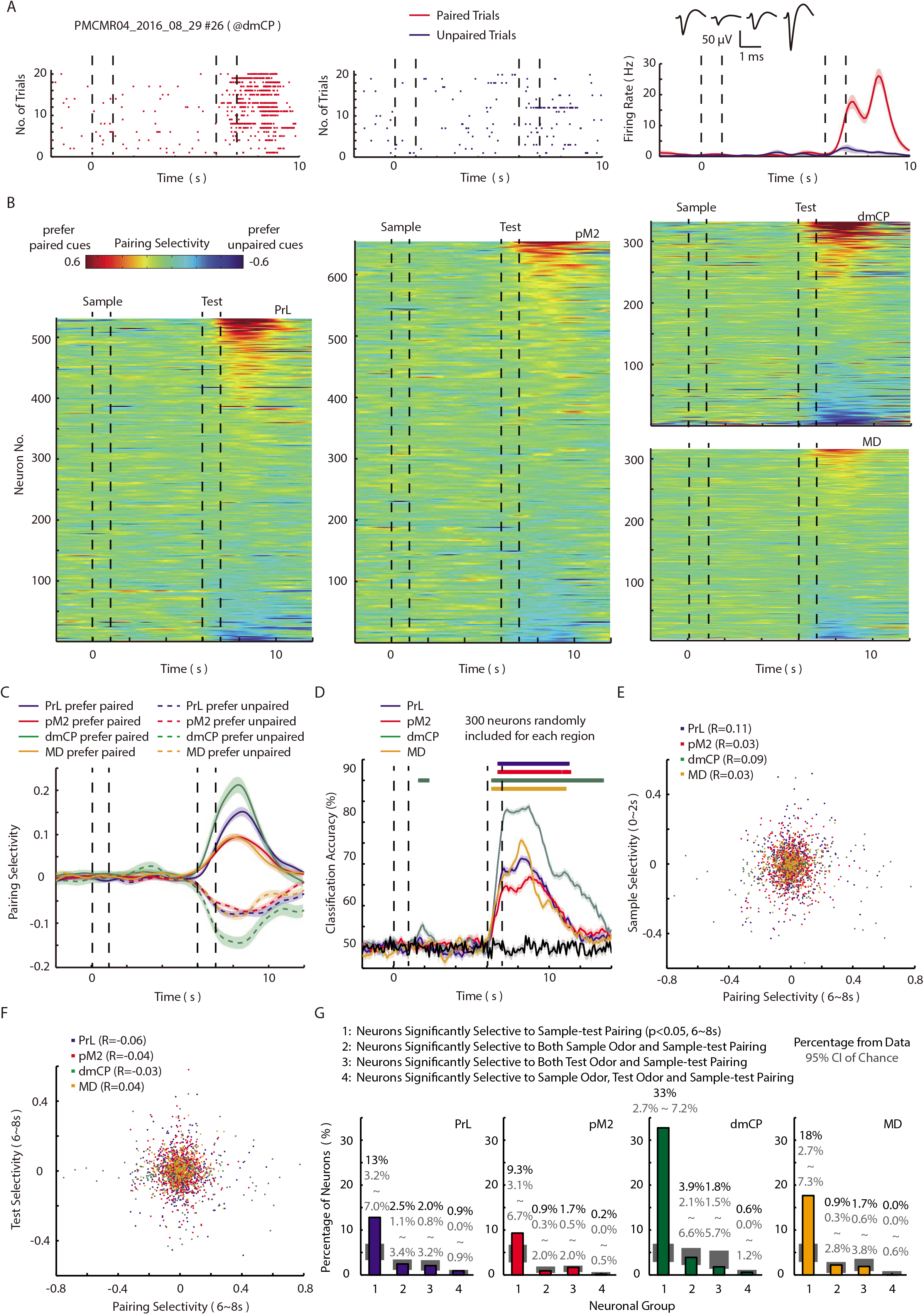
Neurons of the CBT loop represented the pairing-related information. (**A**) A neuron in dmCP with different activity after test-delivery for the trials with paired odors (in red) and non-paired odors (in blue). (**B**) Heat map of pairing selectivity for all neurons recorded in the four regions. Pairing selectivity for each neuron at certain time bin is defined as the difference of average firing rate at this bin from trial with paired sample-test and unpaired sample-test, divided by the summation of them as normalization (Pairing selectivity = (Activty^Paired^ – Activity^Non-paired^) / (Activty^Paired^ + Activity^Non-paired^)). (**C**) Averaged pairing selectivity of neurons preferring paired cues or unpaired cues from PrL, pM2, dmCP and MD. (**D**) Decoding accuracy of sample-test pairing for neurons in PrL, pM2, dmCP and MD. (**F**) Correlation between selectivity for sample-test pairing during 6~8s and selectivity for sample odors during 0~2s for neurons in PrL, pM2, dmCP and MD. (**G**) Correlation between selectivity for sample-test pairing during 6~8s and selectivity for test odors during 6~8s for neurons in PrL, pM2, dmCP and MD. (**G**) Percentage of neurons selective to sample-test pairing, selective to both sample odors and pairing, selective to both test odors and pairing, or all the three information in PrL, pM2, dmCP and MD.

We then tested the relationship between the sensory and pairing coding of the neurons within the CBT loop. Firstly, the correlation between the sample- (**Fig. 3E**) or test-selectivity (**Fig 3F**) and pairing-selectivity were very weak (R^2^ ≤ 0.012), even though some of the correlation was statistically significant (P < 0.05, Spearman correlation, **Fig. 3E-F**). Secondly, the percentage of neurons coding both sensory stimuli and pairing relationship was mostly around chance level (except the neurons of PrL and dmCP, **Fig. 3G**). Therefore, the neurons of the CBT loop tended to code for sensory information and pairing relationship independently. In other words, there were very few paring-selective neurons integrating both sample and test information.

### Further analyses supporting independent coding of sensory information and pairing relationship

To further corroborate the independent coding of sensory information and pairing relationship within the CBT loop, we performed two more independent analyses. Firstly, we performed decoding analysis while removing the neurons coding certain information. The logic is that if a give type of neurons is critical for population decoding, then removing them in decoding analysis should impair the decoding performance. Indeed, when we removed the neurons selective to pairing relationship, there was a marked reduction in decoding performance for pairing relationship (comparing the green with black curves, **Fig. 4A**; statistics in **Fig. 4B**). However, removing the neurons coding for sample and test selective neurons did not change the decoding performance (no difference among black, blue, and red curves, **Fig. 4A**; statistics in **Fig. 4B**). It is noted that the green curves in **Fig. 4A** were still increased one second after the test-odor delivery. This period was corresponding to the water-reward delivery period. The activity during this period was not used to quantify the pairing-selective neurons. Because the water reward was correlated with the hit response in the paired trials, one can infer that there were neurons with reward selectivity during the period, which was indeed shown in SFig. 7. In the current design the coding for water reward cannot be dissociated with the pairing relationship besides the onset timing. Furthermore, the reward prediction cannot be dissociated with the pairing relationship in the paradigm.

**Figure 4:**
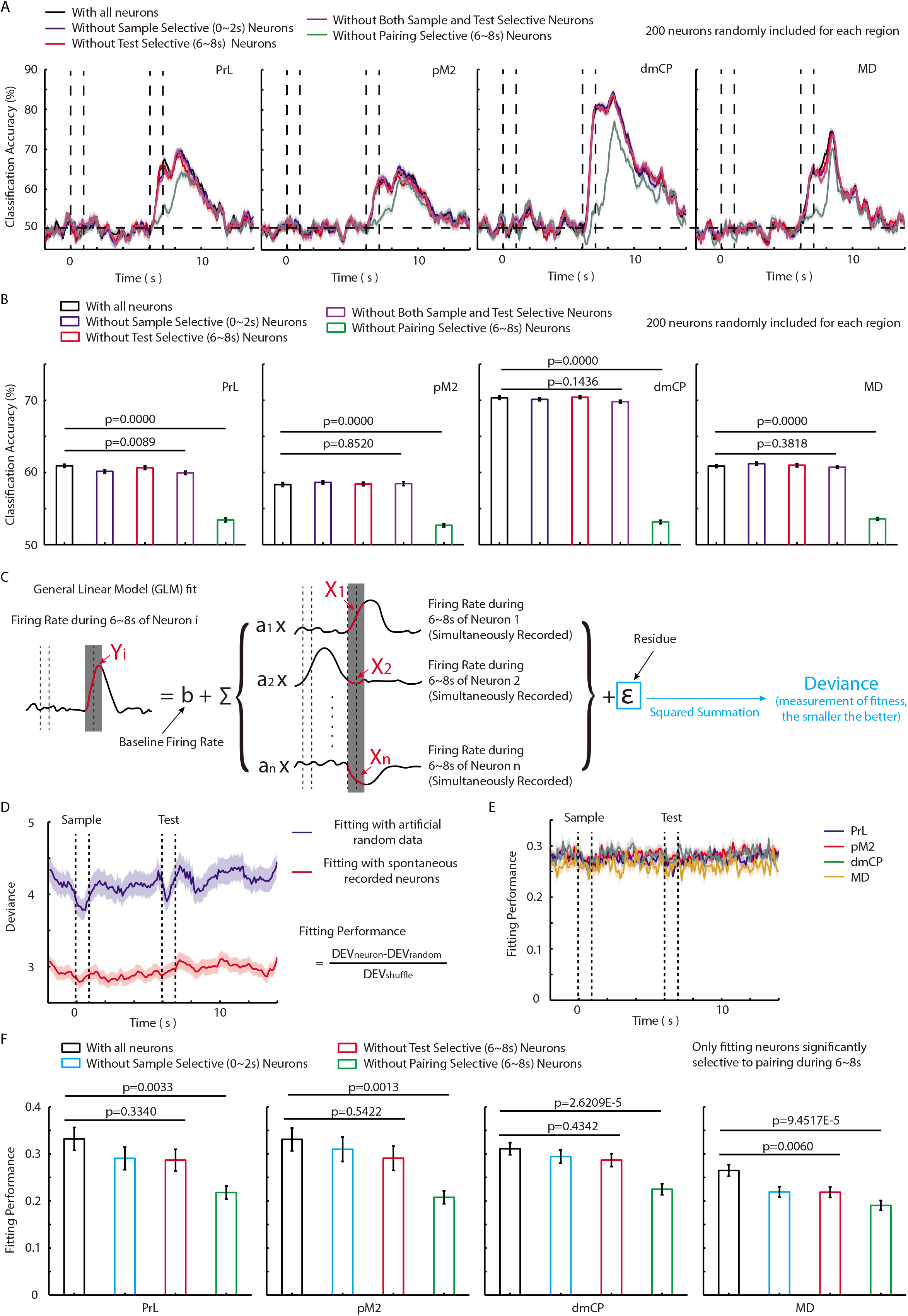
Independent coding of sensory information and pairing relationship. (**A**) Decoding accuracy of sample-test pairing for neurons without sample-selective, test-selective or pairing-selective neurons in PrL, pM2, dmCP and MD. It is noted the significant difference between curves without pairing-selective neurons from other curves. (**B**) The comparing of pairing selectivity of neurons without certain type of neurons in PrL, pM2, dmCP and MD during test odor delivery (6~8s). It is noted that the significant impairment induced by removing pairing-selective neurons. (**C**) Diagram illustrates using generalized linear model to fit the activities of one neuron by other simultaneously recorded neurons. (**D**) The deviance of model using actual recorded neural data is significantly lower than the deviance of model using artificial generated random data. (**E**) Fitting performance for neurons in PrL, pM2, dmCP and MD do not have significant difference and keep stable in the DPA task. Fitting performance is defined by the difference between the deviance of neural model and control model then divide by the deviance of control model. (**F**) The comparing of fitting performance of neurons without certain type of neurons in PrL, pM2, dmCP and MD during test odor delivery (6~8s). It is noted that the significant impairment induced by removing pairing-selective neurons.

Secondly, we fitted the neuronal firing with a general linear model (GLM) to quantify the relative contribution of different types of neurons in explaining the activity of recorded neurons. The task-related variables (including sample/test odors and pairing relationship) and neuronal activities of simultaneously recorded other neurons (from all regions) were incorporated to fit the neuronal activity of each recorded neuron (neural model, **Fig. 4C**). To measure the goodness of fit, deviance is defined as the squared summation of the error between the model prediction and recorded neuronal activity. As a control, we also generated a model incorporating the task-related variables and randomly generated neuronal firing with the same number of simultaneously recorded other neurons (control model). The deviance of control model is significantly larger than that of real-neural model at each time in the task (**Fig. 4D**). Therefore, the neuronal activities can explain firing of other simultaneously recorded neurons, consistent with *(48–50)*. To measure the contribution of a given group of neurons, fitting performance is defined as the difference between the deviance of neural model and the deviance of respective control model normalized by the deviance of control model. The fitting performance keeps stable during the task for PrL, pM2, dmCP and MD (**Fig. 4E**). To identify the contribution of different types neurons in explaining neuronal activity of pairing-selective neurons, we separately calculated the fitting performance without sample-selective, test-selective, or pairing-selective neurons. For PrL, pM2 and dmCP, removing sample-selective or test-selective neurons did not induce significant effects on fitting performance compared with model incorporating all neurons (**Fig. 4F**, blue and red). However, removing pairing-selective neurons significantly impaired the fitting performance (**Fig. 4F**, green). For MD, removing sample-selective or test-selective neurons also significantly impaired the fitting performance. Removing pairing-selective neurons induced significantly stronger impairment. These results supported the dissociation between pairing-selective neurons and sensory-selective neurons.

### Pairing-selective neurons in PrL, pM2 and dmCP are more tightly coupled

The previous GLM was calculated based on the neurons from all regions. To further examine the functional cross-region interaction, we calculated the fitting performance of each region according to model with neurons from only one region in the CBT loop. For example, we fit the PrL neural activity only with firing of pM2 neurons to infer the functional interaction from pM2 neurons to PrL neurons. In other words, such fitting performance indicates the coupling from the neurons of the source region to the neurons in the fitted target region. During either the sample or the test period, the fitting performance among the neurons in PrL, pM2 and dmCP were higher than that of MD (**Fig. 5A** for sample period, **Fig. 5B** for test period). Therefore, PrL, pM2 and dmCP coupled with each other more tightly than they coupled with MD (**Fig. 5C**).

**Figure 5:**
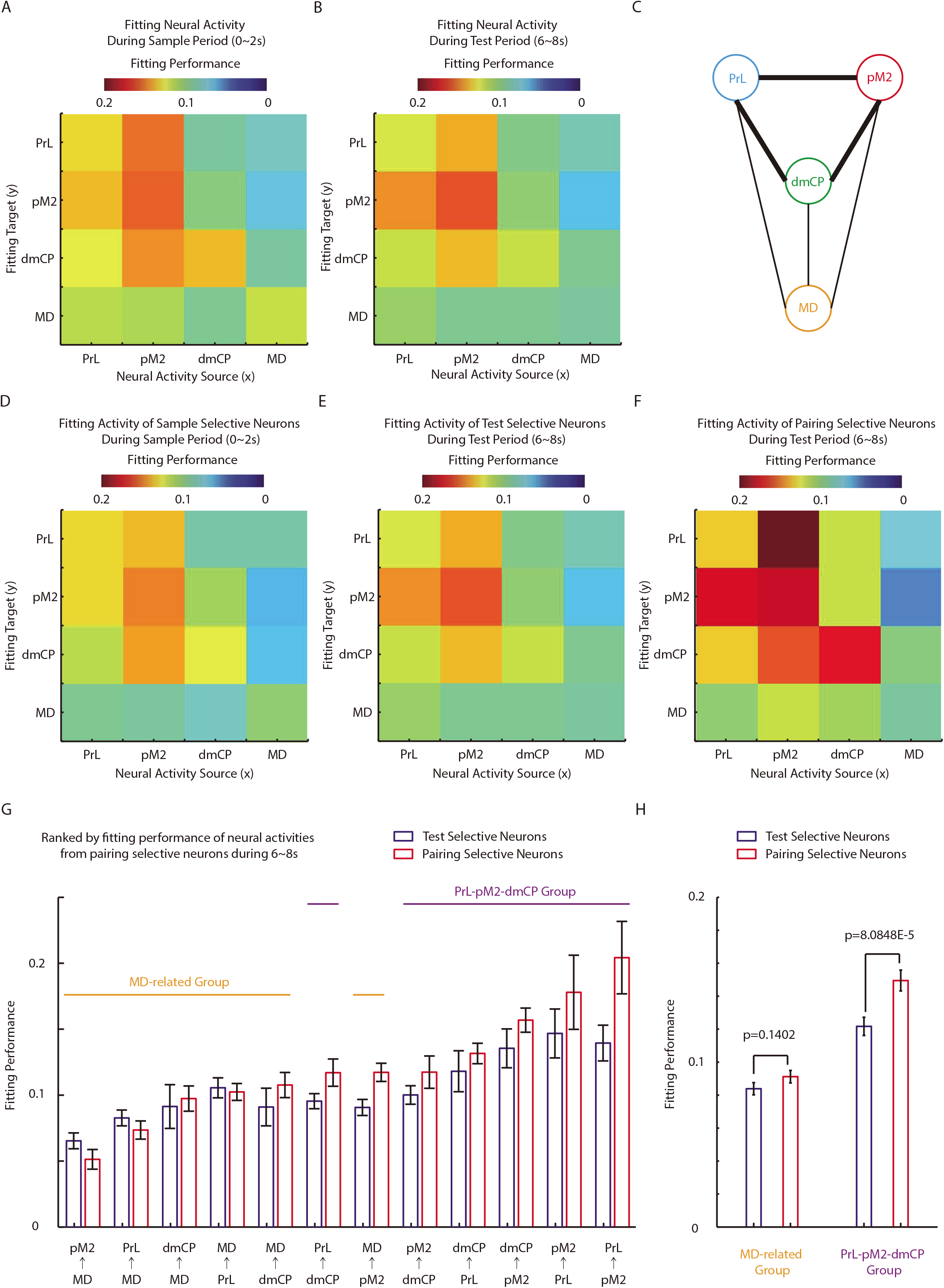
Pairing-selective neurons in PrL, pM2 and dmCP are more tightly coupled. (**A**) Heat-map for the fitting performance of neurons in PrL, pM2, dmCP and MD during sample period. (**B**) Heat-map for the fitting performance of neurons in PrL, pM2, dmCP and MD during test period. (**C**) A diagram illustrating the neural interaction among PrL, pM2, dmCP and MD. (**D**) Heat-map for the fitting performance of sample-selective neurons in PrL, pM2, dmCP and MD during sample period. (**E**) Heat-map for the fitting performance of test-selective neurons in PrL, pM2, dmCP and MD during test period. (**F**) Heat-map for the fitting performance of pairing-selective neurons in PrL, pM2, dmCP and MD during test period. (**G**) The comparing of fitting performance of test-selective neurons and that of pairing-selective neurons in PrL, pM2, dmCP and MD during test period. The rank is according to the fitting performance of pairing-selective neurons. It is noted that most of pairs with low fitting performance is MD related, while most of pairs with high fitting performance is belong to PrL-pM2-dmCP loop. (**H**) The comparing of fitting performance of test-selective neurons and that of pairing-selective neurons in PrL-pM2-dmCP loop and MD-related pairs. It is noted that only neurons in PrL-pM2-dmCP loop significantly increase fitting performance for pairing-selective neurons.

To dissect the functional interaction specific to the coded information, we calculated the fitting performance of regions in the CBT loop only considering the neurons with significant coding for sample, test or pairing information. The coupling pattern calculated for sample-selective neurons (**Fig. 5D**), test-selective neurons (**Fig. 5E**) and pairing-selective neurons (**Fig. 5F**) all had similar patterns as the coupling patterns calculated for all neurons. Specifically, the coupling strength calculated from pairing-selective neurons was stronger than that calculated from sample- or test-selective neurons in the PrL-pM2-dmCP loop (**Fig 5D-F**). This was further demonstrated by arranging the fitting performance of pairing-neurons in all regions (**Fig. 5G**). The ranking for the fitting performance of pairing-neurons is quite similar within PrL-pM2-dmCP loop, which is consistent with the patterns from the heat maps (comparing **Fig. 5G** with **Fig. 5D-F**). We further separated the PrL-pM2-dmCP pairs and MD-related pairs to compare their different coupling patterns. Similarly, we observed higher fitting performance in PrL-pM2-dmCP pairs than MD-related pairs, both for fitting the firing of test-selective and pairing selective neurons (**Fig. 5H**). Furthermore, within the PrL-pM2-dmCP loop, the fitting performance was higher for pairing-selective neurons than test-selective neurons (**Fig. 5H** left, comparing the blue and red bar). This was consistent with the notion that pairing-selective neurons in this loop can be better explained by the firing of other neurons.

### Learning-related dynamics in neuronal firing and functional interaction in the CBT loop

Mice were trained to perform the DPA task well within five days (**Fig. 1C**), therefore enabling us to record and examine the neural correlates during the entire learning process. The fitting performance of pairing-neurons of the PrL-pM2-dmCP pairs was significantly higher on the third day of learning while the fitting performance of pairing-neurons of the MD-related pairs did not have such significant change (**Fig. 6A-B**). We found that sample selectivity was not significantly changed along learning process in PrL, dmCP and MD (**Fig. 6B-C**). We detected a statistically significant change in sample selectivity in pM2 neurons (red curves in **Fig. 6B**), which might be due to the lower selectivity in Day 4 in learning (*post hoc* test). In contrast, the pairing selectivity in all recorded regions was significantly increased during learning in decision-making period (**Fig. 6D**).

**Figure 6:**
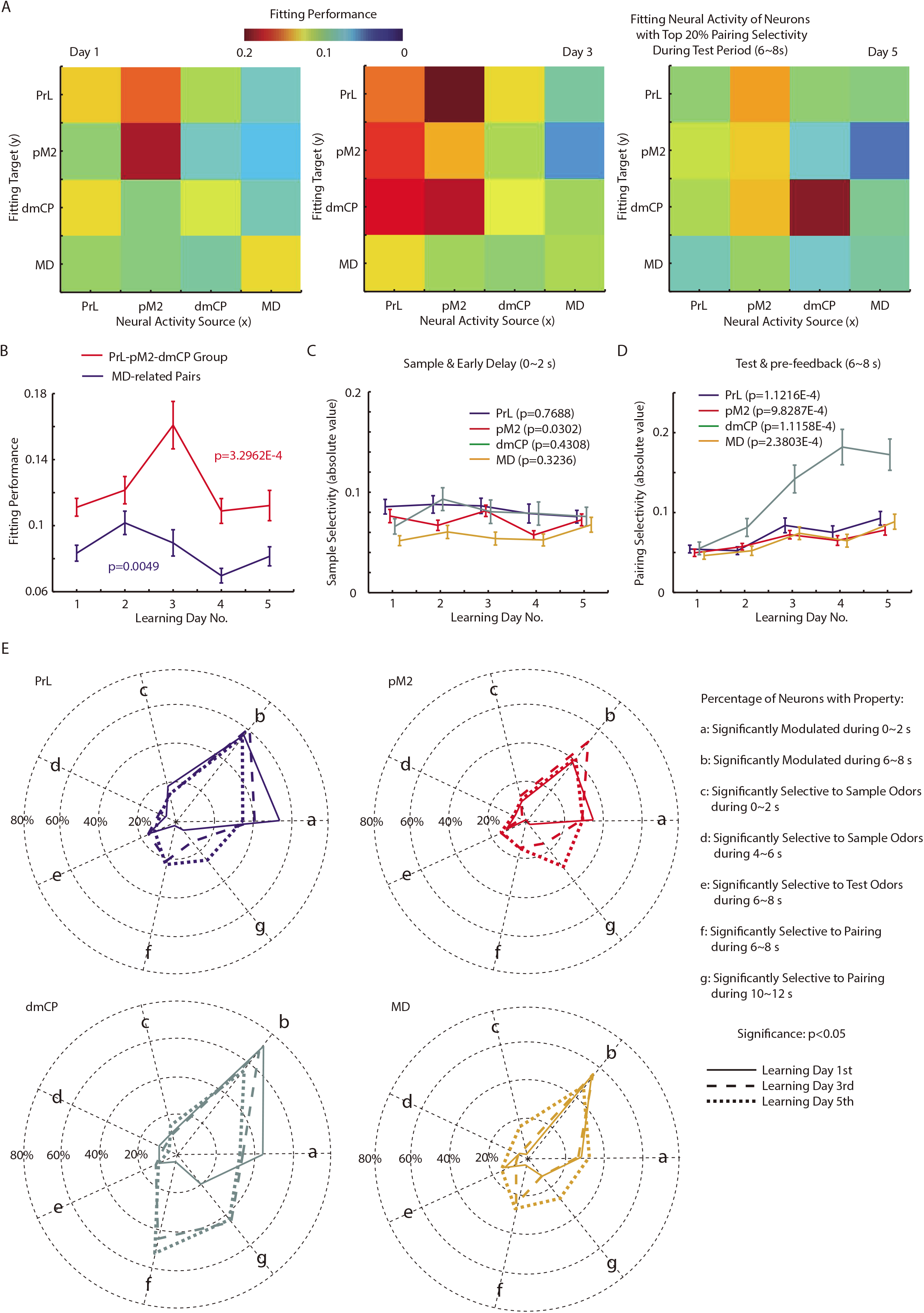
Learning-related dynamics in neuronal firing and functional interaction in the CBT loop. (**A**) Heat-map for the fitting performance of neurons with top 20% pairing selectivity in PrL, pM2, dmCP and MD during test period on the first, third and fifth day of learning the DPA task. (**B**) The fitting performance of neurons with top 20% pairing selectivity in PrL-pM2-dmCP loop and MD-related pairs changed along learning the DPA task. (**C**) Sample selectivity of neurons in PrL, pM2,dmCP and MD keep stable along the learning process. (**D**) Pairing selectivity of neurons in PrL, pM2,dmCP and MD significantly increase along the learning process. (**E**) The percentage of neurons with certain properties in PrL, pM2, dmCP and MD on the first, third and fifth day of learning the DPA task. It is noted that the neural modulation of the four regions decrease, the sensory coding of the four regions keep stable and the decision coding of the four regions increase along the learning process.

To summarize the learning-related dynamics in firing of all the recorded regions, we calculated the percentage of modulated neurons for different time periods on day 1, 3 and 5 of learning process (**Fig. 6F**), including: a) Significantly Modulated during sample-delivery period; b) Significantly modulated during decision-making period; c) Significantly selective to sample odors, during sample-delivery period; d) Significantly selective to sample odors, during late delay period (during 4~6 s after sample odor); e) Significantly selective to test odors during decision-making period; f) Significantly selective to pairing during decision-making period; g) Significantly selective to pairing after reward-feedback (during 10~12 s after sample odor). Generally, the four recorded regions in CBT loop have similar trend of change in firing properties. The neural modulations were slightly decreased along learning (a-b in **Fig. 6F**). The representations for sensory information were stable while the representations for decision were increased along learning (c-g in **Fig. 6F**). Among the four regions, PrL exhibited stronger sensory representation and dmCP exhibited stronger decision representation. Thus, in the recorded CBT loop for the OPDA task, sensory information was stably maintained and the pairing/decision-related information was gradually improved during learning.

## Discussion

In the present study we trained mice to perform a delayed paired association task, in which maintenance of sensory information during delay period and learning association between arbitrary odor pairs are required for a successful task performance. Consistent with the critical role of cortico-subcortical loop in working memory *(18–31)*, all four regions exhibit task-related modulation in neuronal activity (Fig. 2). Interestingly, all four regions also exhibit similar degree of selectivity to sensory information during delay period (Fig. 3). Such a broad distribution in neuronal modulation and selectivity supports the notion of distributed nature of working memory processes *(34)*.

Neuronal activity of all four regions was associated with paired/non-paired relationship during the test-odor delivery and response periods (Fig. 4), with dmCP neurons exhibited the largest difference between the paired *vs*. non-paired trials. The neuronal response during this period can be associated with many brain processes, such as decision making, motor planning, and reward expectation. The exact functional role of neuronal coding of this behavioral phase remains to be determined.

In the current DPA design, mice can either use prospective or retrospective strategy*(46)*. The maintained information in retrospective strategy is the already-presented sample odor, whereas that in prospective strategy is to code for the paired incoming test odor. Our recording study in anterior piriform cortex, a sensory region, suggested that mice were not using prospective strategy to perform the task*(43)*. There was no coding for pairing relationship before test-odor delivery (Fig. 3C), further supporting the idea that mice were not using prospective-coding strategy to perform the task.

When we calculated the percentage of neurons significantly selective for task-related information, the selectivity corresponding to a single stimulus (sample, test and pairing) were higher than chance level (Fig. 2C, 3C). However, the percentages of neurons coding multiple stimuli were close to chance level of overlapping (Fig. 2G, 3G). Therefore, among the four regions, task-related stimulus spread randomly, with low percentage of neurons coding all task-related information. However, the task information must be somehow integrated for optimal SMT. Although the percentage of such neurons is very low for the four regions, we indeed observed a small percentage of them (~1%, Fig. 3G). One hypothesis is that the neurons coding for both sample and test information have very strong projection to other neurons to make them selective for pairing, which need to be determined by future studies.

We generated GLM models to examine the functional interactions among the four regions. The results of the GLM should not be viewed as mono-synaptic connections, because both direct and indirect innervation can contribute to the GLM fitting. However, we could still obtain information about how information functionally flows among the four regions (Citation needed).

In conclusions, we simultaneously recorded multiple neurons from mice PrL, pM2, dmCP and MD. Neurons of the four regions have different level of modulation and selectivity for task-related information. The coding of WM and SMT information is independent. The neurons within PrL-pM2-dmCP loop exhibited stronger coupling than MD, especially for pairing-selective neurons. During learning, the change of firing patterns mostly happens on decision-related neurons and remains stable for sensory-related neurons. Our results provide important insights concerning the dynamic activation and cross-region interaction of a distributed cortico-basal ganglia-thalamus loop in learning a delayed sensorimotor task.

## Acknowledgments

The work was supported by the National Science Foundation for Distinguished Young Scholars of China (31525010, to C.T.L.), the Strategic Priority Research Program of the Chinese Academy of Sciences (Grant No. XDB32010100), the Shanghai Municipal Science and Technology Major Project (Grant No. 2018SHZDZX05), the Instrument Developing Project of the Chinese Academy of Sciences (Grant No. YZ201540), the Key Project of Shanghai Science and Technology Commission (No.15JC1400102, 16JC1400101), the General Program of Chinese National Science Foundation (31471049), the China–Netherlands CAS-NWO Programme: The Future of Brain and Cognition (153D31KYSB20160106).

## Reference

1. G. E. Alexander, M. R. DeLong, P. L. Strick, Parallel organization of functionally segregated circuits linking basal ganglia and cortex. Annu Rev Neurosci 9, 357–381 (1986).

2. A. M. Graybiel, T. Aosaki, A. W. Flaherty, M. Kimura, The basal ganglia and adaptive motor control. Science 265, 1826–1831 (1994).

3. O. Hikosaka, Y. Takikawa, R. Kawagoe, Role of the basal ganglia in the control of purposive saccadic eye movements. Physiol Rev 80, 953–978 (2000).

4. S. Grillner, J. Hellgren, A. Menard, K. Saitoh, M. A. Wikstrom, Mechanisms for selection of basic motor programs--roles for the striatum and pallidum. Trends Neurosci 28, 364–370 (2005).

5. M. Jahanshahi, I. Obeso, J. C. Rothwell, J. A. Obeso, A fronto-striato-subthalamic-pallidal network for goal-directed and habitual inhibition. Nat Rev Neurosci 16, 719–732 (2015).

6. X. Jin, R. M. Costa, Shaping action sequences in basal ganglia circuits. Current opinion in neurobiology 33, 188–196 (2015).

7. L. A. Gunaydin, A. C. Kreitzer, Cortico-Basal Ganglia Circuit Function in Psychiatric Disease. Annu Rev Physiol 78, 327–350 (2016).

8. E. A. Murray, S. P. Wise, K. S. Graham, The evolution of memory systems : ancestors, anatomy, and adaptations. (Oxford University Press, Oxford, United Kingdom; New York, NY, ed. First Edition., 2017), pp. xvii, 496 pages.

9. A. V. Kravitz et al., Regulation of parkinsonian motor behaviours by optogenetic control of basal ganglia circuitry. Nature 466, 622–626 (2010).

10. G. Cui et al., Concurrent activation of striatal direct and indirect pathways during action initiation. Nature 494, 238–242 (2013).

11. L. H. Tai, A. M. Lee, N. Benavidez, A. Bonci, L. Wilbrecht, Transient stimulation of distinct subpopulations of striatal neurons mimics changes in action value. Nature neuroscience 15, 1281–1289 (2012).

12. A. V. Kravitz, L. D. Tye, A. C. Kreitzer, Distinct roles for direct and indirect pathway striatal neurons in reinforcement. Nature neuroscience 15, 816–818 (2012).

13. M. K. Lobo et al., Cell type-specific loss of BDNF signaling mimics optogenetic control of cocaine reward. Science 330, 385–390 (2010).

14. L. Ding, J. I. Gold, Caudate encodes multiple computations for perceptual decisions. The Journal of neuroscience : the official journal of the Society for Neuroscience 30, 15747–15759 (2010).

15. B. U. Forstmann et al., Cortico-striatal connections predict control over speed and accuracy in perceptual decision making. Proceedings of the National Academy of Sciences of the United States of America 107, 15916–15920 (2010).

16. J. Grinband, J. Hirsch, V. P. Ferrera, A neural representation of categorization uncertainty in the human brain. Neuron 49, 757–763 (2006).

17. A. R. Aron et al., Inhibition of subliminally primed responses is mediated by the caudate and thalamus: evidence from functional MRI and Huntington’s disease. Brain 126, 713–723 (2003).

18. A. Isseroff, H. E. Rosvold, T. W. Galkin, P. S. Goldman-Rakic, Spatial memory impairments following damage to the mediodorsal nucleus of the thalamus in rhesus monkeys. Brain Res 232, 97–113 (1982).

19. J. M. Fuster, The prefrontal cortex : anatomy, physiology, and neuropsychology of the frontal lobe. (Lippincott-Raven, Philadelphia, ed. 3rd, 1997), pp. xvi, 333 p.

20. S. B. Floresco, D. N. Braaksma, A. G. Phillips, Thalamic-cortical-striatal circuitry subserves working memory during delayed responding on a radial arm maze. The Journal of neuroscience : the official journal of the Society for Neuroscience 19, 11061–11071 (1999).

21. A. Hernandez et al., Decoding a perceptual decision process across cortex. Neuron 66, 300–314 (2010).

22. T. D. Barnes et al., Advance cueing produces enhanced action-boundary patterns of spike activity in the sensorimotor striatum. Journal of neurophysiology 105, 1861–1878 (2011).

23. K. L. Mills et al., Altered cortico-striatal-thalamic connectivity in relation to spatial working memory capacity in children with ADHD. Front Psychiatry 3, 2 (2012).

24. Y Watanabe, S. Funahashi, Thalamic mediodorsal nucleus and working memory. Neurosci Biobehav Rev 36, 134–142 (2012).

25. S. Parnaudeau et al., Inhibition of mediodorsal thalamus disrupts thalamofrontal connectivity and cognition. Neuron 77, 1151–1162 (2013).

26. Z. V. Guo et al., Flow of cortical activity underlying a tactile decision in mice. Neuron 81, 179–194 (2014).

27. D. Liu et al., Medial prefrontal activity during delay period contributes to learning of a working memory task. Science 346, 458–463 (2014).

28. N. Li, T. W. Chen, Z. V. Guo, C. R. Gerfen, K. Svoboda, A motor cortex circuit for motor planning and movement. Nature 519, 51–56 (2015).

29. S. S. Bolkan et al., Thalamic projections sustain prefrontal activity during working memory maintenance. Nature neuroscience 20, 987–996 (2017).

30. T. Kamigaki, Y. Dan, Delay activity of specific prefrontal interneuron subtypes modulates memory-guided behavior. Nature neuroscience, (2017).

31. L. I. Schmitt et al., Thalamic amplification of cortical connectivity sustains attentional control. Nature 545, 219–223 (2017).

32. J. Jonides et al., The mind and brain of short-term memory. Annual review of psychology 59, 193–224 (2008).

33. A. Baddeley, Working memory: theories, models, and controversies. Annual review of psychology 63, 1–29 (2012).

34. T. B. Christophel, P. C. Klink, B. Spitzer, P. R. Roelfsema, J. D. Haynes, The Distributed Nature of Working Memory. Trends Cogn Sci 21, 111–124 (2017).

35. M. N. Shadlen, W. T. Newsome, Neural basis of a perceptual decision in the parietal cortex (area LIP) of the rhesus monkey. Journal of neurophysiology 86, 1916–1936 (2001).

36. P. Cisek, J. F. Kalaska, Neural correlates of reaching decisions in dorsal premotor cortex: specification of multiple direction choices and final selection of action. Neuron 45, 801–814 (2005).

37. M. L. Platt, P. W. Glimcher, Responses of intraparietal neurons to saccadic targets and visual distractors. Journal of neurophysiology 78, 1574–1589 (1997).

38. S. Crochet, S. H. Lee, C. C. H. Petersen, Neural Circuits for Goal-Directed Sensorimotor Transformations. Trends Neurosci 42, 66–77 (2019).

39. E. H. Baeg et al., Dynamics of population code for working memory in the prefrontal cortex. Neuron 40, 177–188 (2003).

40. X. L. Qi, T. Meyer, T. R. Stanford, C. Constantinidis, Changes in prefrontal neuronal activity after learning to perform a spatial working memory task. Cereb Cortex 21, 2722–2732 (2011).

41. T. A. Engel, X. J. Wang, Same or different? A neural circuit mechanism of similarity-based pattern match decision making. The Journal of neuroscience : the official journal of the Society for Neuroscience 31, 6982–6996 (2011).

42. Z. Han, X. Zhang, J. Zhu, Y Chen, C. T. Li, High-Throughput Automatic Training System for Odor-Based Learned Behaviors in Head-Fixed Mice. Front Neural Circuits 12, 15 (2018).

43. X. Zhang et al., Active information maintenance in working memory by a sensory cortex. bioRxiv, 385393 (2018).

44. R. Romo, C. D. Brody, A. Hernandez, L. Lemus, Neuronal correlates of parametric working memory in the prefrontal cortex. Nature 399, 470–473 (1999).

45. S. Funahashi, C. J. Bruce, P. S. Goldman-Rakic, Mnemonic coding of visual space in the monkey’s dorsolateral prefrontal cortex. Journal of neurophysiology 61, 331–349 (1989).

46. G. Rainer, S. C. Rao, E. K. Miller, Prospective coding for objects in primate prefrontal cortex. The Journal of neuroscience : the official journal of the Society for Neuroscience 19, 5493–5505 (1999).

47. E. M. Meyers, D. J. Freedman, G. Kreiman, E. K. Miller, T. Poggio, Dynamic population coding of category information in inferior temporal and prefrontal cortex. Journal of neurophysiology 100, 1407–1419 (2008).

48. C. A. Runyan, E. Piasini, S. Panzeri, C. D. Harvey, Distinct timescales of population coding across cortex. Nature 548, 92–96 (2017).

49. R. V. Rikhye, A. Gilra, M. M. Halassa, Thalamic regulation of switching between cortical representations enables cognitive flexibility. Nature neuroscience 21, 1753–1763 (2018).

50. E. J. Hwang, J. E. Dahlen, M. Mukundan, T. Komiyama, History-based action selection bias in posterior parietal cortex. Nat Commun 8, 1242 (2017).

