## Supplementary material for "Dynamic neuronal activation of a distributed cortico-basal ganglia-thalamus loop in learning a delayed sensorimotor task": Materials and methods

**Material and Methods**

**Animals**

Male adult C57BL/6 mice (SLAC, as wild type) were used for the current study (8-12 weeks of age, weighted between 20 to 30 g). Wild type mice were provided by the Shanghai Laboratory Animal Center (SLAC), CAS, Shanghai, China. Mice were group-housed (4-6/cage) under a 12-h light-dark cycle (light on from 6:30 a.m. to 6:30 p.m.). Before behavioral training, mice were housed in stable conditions with food and water *ad libitum*. After the start of behavioral training, water supply was restricted. Mice could drink water only during and immediately after training. Care was taken to keep mice body weight (b.w.) above 80% of normal level. All animal studies and experimental procedures were approved by the Animal Care and Use Committee of the Institute of Neuroscience, Chinese Academy of Sciences, Shanghai, China.

**Behavioral setups**

We utilized an olfaction based delayed pair-associative (DPA) task in head-fixed mice. Computer controlled olfactometry systems were used for semi-automatic training. Microprocessor based controller was used to switch on/off the solenoid valves and peristaltic pump for controlling water and odor delivery in millisecond temporal resolution. Three-way solenoid valves were used for controlling air flow, whereas peristaltic pump were used for controlling water flow. Total length of odor-delivery tubes (inner diameter: 2.5 mm; outer diameter: 4.0 mm) was minimized to increase the turnover rate of odorants. Butyl formate (S1), 1-pentanone (S2), ethyl acetate (T1), methyl butenol (T2) were used at concentrations of 1:500 (v/v in mineral oil, O122-4). We measured the concentration of odor by photoionization detector (PID). The readout of PID during the delay period was similar to that of the baseline level, indicating for efficient clearance of residual odor. In hit trials water was provided ~10 μl water for 50 msec in a response time window. Multiple behavioral setups (6 for behavior training, 2 for extracellular recording) were used. Independent controller was used for each behavioral setup. Odor and water supply were independent for all behavioral setups. All facility parameters, including length for all tubes and air-flow rate, were kept the same across all behavioral setups. Behavioral results from 2 or 6 controllers were recorded simultaneously by custom written software and stored in computers.

**Behavior Training**

In DPA task, a sample olfactory stimulus was presented at the start of a trial, followed by a delay-period (5 sec) and then a testing stimulus. Odor delivery duration was set to one second, which was sufficient for mice to perceive olfactory cues. Mice were trained to lick in the response window. The response window (0.5 sec in duration) was started 1 sec after the offset of the second odor delivery. Licking events detected in the response window in paired trials were regarded as ‘Hit’ and will trigger instantaneous water delivery (50 ms in duration). ‘False alarm’ was defined by the detection of licking events in the response window in non-paired trials and mice were not punished in false alarm trials. Mice were neither punished nor rewarded for ‘Miss’ (no-lick in a paired trial) nor ‘Correct Rejection’ (CR, no-lick in a non-paired trial) trials. Licking events were detected by infrared beam breakers. Odor and water delivery, laser illumination, and licking events were recorded by computers through serial ports. In each day, mice were required to perform 240 trials for DPA task in optogenetic and electrophysiological experiments. After training sessions ended each day, mice were supplied with free water until satiety.

Before the start of training, mice were water restricted for 2 days. The behavioral training process included habituation, shaping and task learning phases. In habituation phase, mice were head-fixed in behavioral setups and trained to lick water from a water tube, encouraged with automatically delivered water through rodent lavage needles. Typically in 1 to 2 days, mice could learn to lick for 1 to 2 minutes continuously. The shaping phase was then started, in which only paired trials were applied and water was provided in all trials each day. In the beginning of shaping phase, water was delivered manually through syringes to encourage mice to lick in the response window for only 10 trials. For the rest trials, mice were trained automatically. Typically the shaping phase lasted for 2 to 3 days.

The task learning phase was then started from the next day, which was defined as day 1 in the behavioral analysis reported in all figures. All kinds of trials were applied pseudo-randomly, i.e., two paired and two unpaired trials of balanced odor-pairs were presented randomly in every consecutive four trials in DPA task. No human intervene was applied in task learning phase to minimize any potential human bias in behavioral results. The performance correct rate (referred to as “performance” in labels of figures) of each session was defined by:

*Performance correct rate = (num. hit trials + num. correct rejection trials) / total number of trials*

*Hit*, *False choice*, and *Correct rejection rates* were defined as follows:

*Hit rate = num. hit trials / (num. hit trials + num. miss trials)*

*False choice rate =* *num. false choice trials / (num. false choice trials + num. correct rejection trials)*

*Correct rejection rate* = *num. correct rejection trials/ (num. false choice trials + num. correct rejection trials)*

**Assembly of chamber for microdrive**

A chamber of a microdrive was composed of eight pieces of printed circuit boards (PCB) (0.05 mm thickness), which were designed with Altium Designer and manufactured in Binqidianzi, Shanghai, China. Two pieces of PCBs were stacked and glued together to form the ceiling of a microdrive. Holes on the ceiling were for holding the larger end of screws (11 mm in length and 2.0 mm in diameter). Two pieces of PCBs were stacked and glued together to form the floor of a microdrive. Two smaller holes on the floor were for restricting the smaller end of screws. Other sets of smaller holes were used to hold two metal rods, which was designed to prevent nuts from rotating while turning screw head. This design transferred rotation of screw heads into movement of nuts through screw axis, therefore drove tetrodes into brain tissue. Each microdrive had two independently movable screws and corresponding nuts. Two pieces of PCBs were glued to be side plates of the chamber.

**Assembly of tetrodes**

Each tetrode was constructed with polyimide insulated, Ni-Chrome wire (12.5 μm core diameter for tetrode). Construction of a tetrode was started by obtaining a 20 cm long wire. Wire was folded in half for twice, and then its open end was clamped together with a clip. The connected end of wires was hanged onto a horizontal bar. A rotating force was applied to the clip manually to trigger counter-clock wise twists. After the counterclockwise twisting stopped, freely clockwise unwinding was allowed until the clip stopped rotating. After tetrode twisting was completed, insulation coats of wires were melted and fused together by gently heating with a heat gun (200° C) for 4-5 sec. Tetrode was removed from the twisting apparatus by gently lifting the clip and cutting the wire near the clip. The clip was gently lifted and the tetrode wire was cut near the clip. Tetrode then was removed from the twisting apparatus. At the connected end of wires, the loop was cut into four non-bonded strands of equal length. Individual strands were separated by gently bending wires with a soft tipped tweezer. Insulation coats of strand tips were removed carefully. Then wire tips were soldered to corresponding pins on a PCB connector which was electrically connected to an adaptor. Two electrodes for ground and reference (magnet wire, 0.01 mm^2^) were soldered to the corresponding pins. Each pin was individually coated with silver paint to enhance conductance. Connector pin arrays and Omnetics adaptors were then coated with silica gel to protect connection between wires and pins

**Assembly of microdrive**

Pieces of polyimide tubing (inner diameter 75 μm, outer diameter 150 μm) were glued together (2 × 4) to form guide tubes. They were then inserted and glued on the wall of lower holes. Tetrode wires were inserted into guide tubes from the side of a micro-drive chamber. Then with epoxy glue (5 min epoxy system), middle part of tetrode wires were fixed on each of two independently movable screw nuts in a microdrive chamber. After epoxy glue dried, electrode end of tetrodes were trimmed and the length out of guide tubes was adjusted. Trimmed tips of tetrodes were electroplated to a final impedance of about 1MΩ at 1kHz, with an automatic multichannel electroplating system. In electroplating, low concentration of gold solution was prepared by mixing with solution of polyethylene glycol (PEG, 1mg/mL, v/v) at a concentration of 1:9 (Gold : PEG). Omnetics adaptors were then fixed on side plates of a microdrive chamber.

**Surgical implantation of tetrode micro-drive**

The implantation procedure was similar to that for implantation of optical fibers. A micro-drive was sterilized by ultraviolet radiation for more than 20 minutes. Several cranial windows of 1 by 1 mm were made in each hemisphere for implantation. The center of an electrode array was targeted to AP +2.00 mm, ML 0.40 mm, and DV 1.65 mm for PrL, AP +0.86 mm, ML 0.50 mm, and DV 0.50 mm for pM2, AP +0.86 mm, ML 1.20 mm, and DV 2.50 mm for dmCP, AP -1.22 mm, ML 0.50 mm, and DV 2.90 mm for MD. Dura mater was carefully removed with surgery needles with as less bleeding as possible. Similar to optical fiber implantation, tissue gel (3M, US) and dental cement were carefully applied without relative motion between a micro-drive and brain tissue. Antibiotic drug (ampicillin sodium, 20 mg/mL, 160 mg/kg b.w.) was injected for three consecutive days after surgery.

**Electrophysiological Recording**

After surgery, mice were allowed to recover for at least one week before behavioral training. The recording began from the end of shaping period. We did not select cells to be recorded to ensure independent sampling. Wide band signals (0.5-8000 Hz) from all tetrodes were amplified (× 20000) and digitized at 40 kHz with the Multi-neuron Acquisition Processor and all data were saved to a hard-disk. Spike event detection and sorting was performed offline as described below. Raw signal was filtered online (250 - 8000 Hz) and only spikes detected across threshold (5 folds of STD) were stored for further analysis.

**Analysis: Spike sorting**

Care was taken to ensure that only single-units were sorted and analyzed, based on clustering analysis on principal components (PC) of spike waveforms. Offline spike detection was performed with OfflineSorter. Raw signals were filtered in 250~8000 Hz to remove field potentials. Typically negative six times of standard deviation of recorded signals of each lid of a tetrode were set as thresholds for detecting spike events. Deflections lower than the threshold were marked as putative spike events. Spike events that were detected at any lead of a tetrode would retrieve corresponding waveforms at all four lead for further analysis. PCA was performed for tetrode-waveforms to extract the first three PCs explaining the largest variance. Then, “T-Dist EM” clustering provided by OfflineSorter was performed in 3D PC space of waveforms. Single neuron was included only if there were no more than 0.1% of spikes within 2 ms refractory period and the averaged firing rate was higher than 2 Hz. Recording stability was verified by visually inspection of PCs projection of spike waveforms throughout recording. To ensure genuine single-units, we plotted and inspected spike amplitudes and peak-to-valley intervals of recorded spikes. In this plot, multiple clusters meant multiple neurons or a noise source were included. A sharp cut-off in either side of distribution meant significant missed spike events. We therefore excluded the units from the further analysis in either of above cases. Including them in the analysis, however, did not affect the results in learning specific modulation in delay-period activities, PCA trajectories, decoding, and correlation with behavioral performance (data not shown). To guarantee independent sampling of neurons across different days, we compared spike waveforms and autocorrelation to detect putative same neurons recorded in different days. For those neurons, only the activity of the first recorded day was included.

**Analysis: Normalized firing rate and activity heat map**

All further analyses were performed by custom-written codes with MATLAB. Statistical significance was defined as p < 0.05 unless noted otherwise. Baseline period was defined as -2 ~ 0 second before onset of odor sample. Firing rate from each trial of baseline was averaged to form baseline activity vector of each neurons. Mean and standard deviation of this baseline activity vector were used to convert averaged firing rate of different time bins (size: 100 ms) into Z-score. Activity of all neurons were sorted by the mean Z-scored firing rate and plotted as heat map using ‘Jet’ color-map defined in Matlab. To quantify the difference of modulation in different event periods, we used absolute value of normalized firing rate as normalized firing modulation. We compared the normalized firing modulation during task event period with that during baseline to measure firing rate change induced by task stimulus.

To keep the time duration of neural activities same across different event periods, the time window of ‘sample period’ in neural activity analysis was defined as 0~2 second of the timeline (including 1 second of sample odor duration and 1second after the offset of sample odor). The time window of ‘test period’ in neural activity analysis was defined as 6~8 second of the timeline (including 1 second of test odor duration and 1second after the offset of test odor). The time window of ‘late delay period’ in neural activity analysis was defined as 4~6 second of the timeline (from 3 seconds after the offset of sample odor to the onset of test odor).

**Analysis: Selectivity**

Neuronal selectivity for certain binary stimulus was defined as:

$$Selectivity=\frac{{Firing Rate}_{stimulus 1}-{Firing Rate}_{stimulus 2}}{{Firing Rate}_{stimulus 1}+{Firing Rate}_{stimulus 2}}$$

Here the firing rate was the average value across all trials on each 100ms bin. Selectivity of all neurons were sorted by the mean selectivity and plotted as heat map using ‘Jet’ color-map defined in Matlab. To quantify the difference of selectivity in different event periods, we used absolute value of selectivity as absolute selectivity. We compared the absolute selectivity during task event period with that during baseline to measure neuronal coding power change induced by task stimulus.

To calculate neuronal selectivity for sample, stimulus 1 is S1 and stimulus 2 is S2;

To calculate neuronal selectivity for test, stimulus 1 isT1 and stimulus 2 is T2;

To calculate neuronal selectivity for pairing, stimulus 1 is paired condition and stimulus 2 is unpaired condition;

**Analysis: Neuron percentage with significant firing property**

Neurons with certain property (significant licking modulation, significant test odor selectivity, etc.) were defined by neurons with significant different firing rate (Mann-Whitney U-test, p<0.05) during interested window in trials under two conditions (trials with S1 or S2, trials with paired cues or unpaired cues, etc.). Percentage of such neurons was the ratio between number of such neurons and all neurons.

**Analysis: Decoding**

We perform the decoding analysis based on the classifier of maximum correlation coefficient (MCC):

1) Because the numbers of the recorded neurons were different for PrL, pM2, dmCP and MD, 300 neurons were randomly selected for each region. This allowed a fair comparison in decoding efficiency.
2) We randomly selected 40 trials for each condition (e.g. trails with S1 or S2). The duration of the whole trial was 18.7 sec. The bin size for calculating the firing rate was 100 ms and 187 bins were created for each trial. The resulted population activity matrix had the dimension of 300 X 40 X 187.
3) For each condition, the activity template was created by averaging firing rate from half of the trials (20 randomly selected out of 40) for all the neurons. The resulting template had the dimension of 300 X 187.
4) The rest 20 trials for given condition was selected for all 300 neurons as data for test, resulting in 20 testing matrixes with the dimension of 300 X 187.
5) In testing phase, for each time point, we generated two testing vectors (with the dimension of 300 X 1) and two template vectors (each with the dimension of 300 X 1), corresponding to each condition.
6) We separately calculated the correlation coefficients between each of the testing vectors and each of the two template vectors.
7) The two testing vectors were assigned as condition1 or condition2 according the larger correlation coefficient to the corresponding template vector.
8) The truth score of 1 or 0 was assigned if the assignment in 7^th^ step was correct or not, respectively.
9) The same procedure from 5^th^ to 8^th^ was repeated for all the time bins, to generate a truth score vector (with the dimension of 2 X 187).
10) The procedure from 5^th^ to 9^th^ was repeated for all 20 test matrixes. The resulting 40 X 187 matrix was averaged to obtain a classification accuracy vector (with the dimension of 1 X 187).
11) The procedure from 5^th^ to 10^th^ was repeated for 50 times to obtain a 50 X 187 classification accuracy matrix. The means (1 X 187) and SEMs (1 X 187) of this matrix were used to plot figures.

To generate the shuffled data, the IDs of conditions and neurons from all regions were randomly re-assigned in step 3 to generate a shuffled template. Other procedures were followed as previously to generate the means and SEMs of the resulted classification accuracy matrix for the shuffled data.

**Analysis: Generalized Linear Model**

We performed generalized liner model to fit firing counts of each neuron in each 100ms bin during DPA task by simultaneously recorded other neurons. The fitting was performed by Matlab with function ‘glmfit’.

1) For each mouse in one learning day for DPA task, 80 trials were picked for analysis. For all neurons recorded (neuron number is ‘N’), the firing counts in each 100ms bin was calculated. Because the duration of each trial is 18.7s, the data set is one N X 187 X 80 matrix.
2) For each neuron at certain time bin, the fitting target is the firing counts in this time bin across trials, which is one 80 X 1 vector (vector ‘y’). The first part of the fitting source was the behavior parameter of each trial, including sample odor, test odor and pairing condition, which is one 80 X 3 matrix. The second part of the fitting source is the firing counts of the remaining (N-1) neurons, which is one 80 X (N-1) matrix. Therefore, the total fitting source is ones 80 X (N+2) matrix (matrix ‘X’).
3) A weight vector ‘a’ and a constant ‘b’ was calculated by function ‘glmfit’ in matlab to make ‘Xa+b’ as close to ‘y’ as possible. The residue of this fitting is ε=y-(Xa+b). As a measure, the deviance ‘DEVneural’ is defined as the squared summation of the residue ε.
4) To compare the performance of the model consisted with simultaneously recorded neural activities, control model using artificially generated random matrix with same dimension of 80 X (N-1) as the second part of fitting source. Similar as the calculation in 3), ‘DEVrandom’ is calculated respectively. The fitting performance of the behavior parameters and simultaneously recorded neurons is defined as **|**DEVneural- DEVrandom**|/** DEVrandom, which indicate how better neural data can do than artificial random data.
5) Therefore, for each neuron in each time bin, there is one fitting performance. The fitting performance of all recorded neurons for the mouse in this learning day is one N X 187 X 80 matrix. We average the fitting performance across all trials to get one N X 187 matrix, which is used to plot figures.
6) To identify the importance of neurons with certain properties (sample selectivity, pairing selectivity, etc.), we remove neurons with certain properties in 2) to detect the impairment on the final fitting performance. The decrease of fitting performance indicates the importance of removed neurons in fitting. In these situations, for control model, the artificially generated random data always keep the same dimension as the neural data.
7) To measure the coupling between neurons from different regions, we only use neurons from one region (PrL, pM2, dmCP and MD) in 2) and calculate the final fitting performance. Higher fitting performance in this situation means better explanation to the target neuron by neurons in this region.

Fitting performance of each neuron would be grouped according to different region, different neuron types (sample selectivity, pairing selectivity, etc.) or in different learning day to detect the unique properties of certain group of neurons.
