## Supplementary figures for "Dynamic neuronal activation of a distributed cortico-basal ganglia-thalamus loop in learning a delayed sensorimotor task"

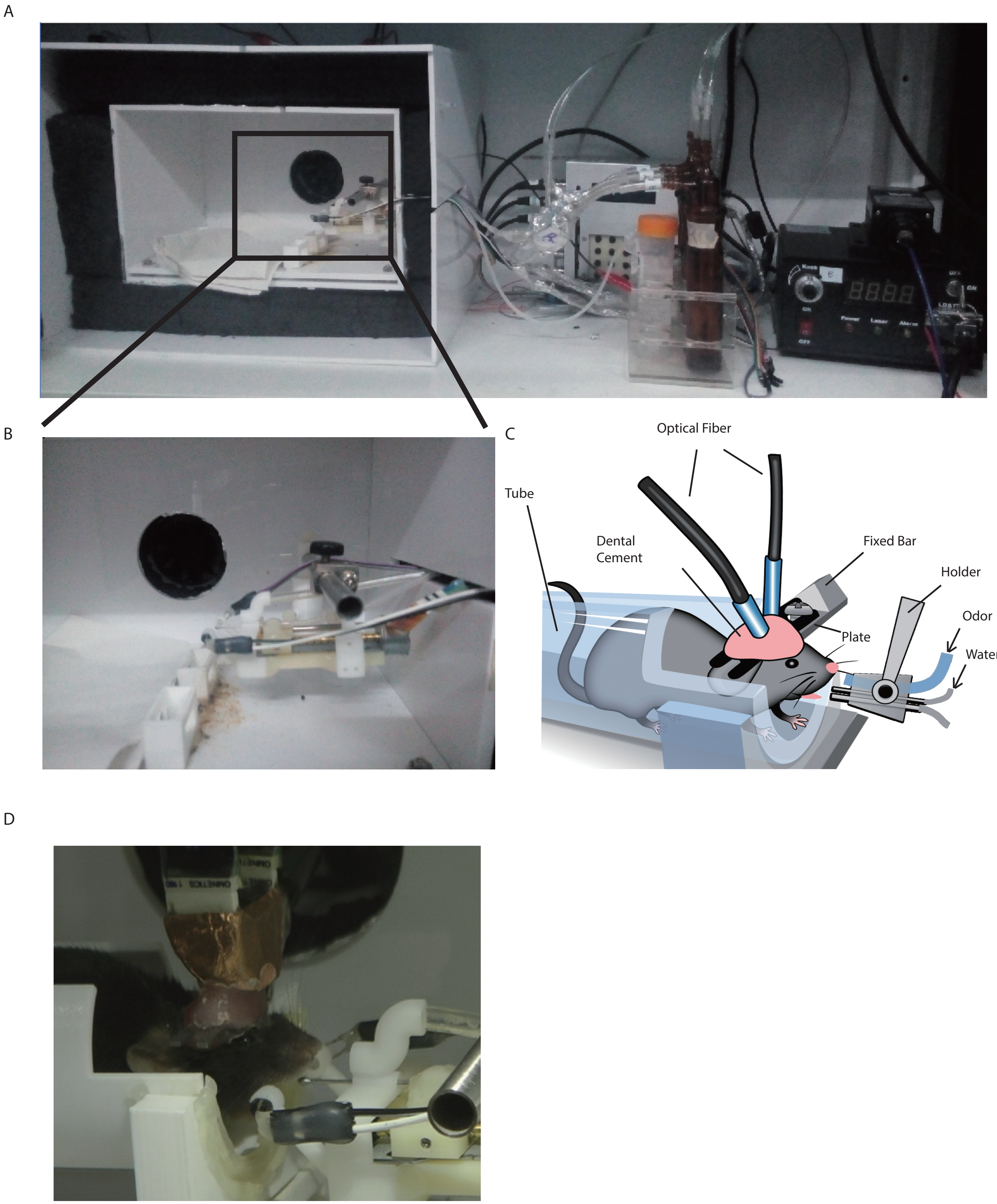


Supplementary Fig. 1. Training system of olfactory memory task on mice. (A) A photo of the whole training system including training box, controller, odorants, etc. (B) Enlarged photo of the black frame in (A). (C) A diagram for the structures in (B) with a mouse. (D) A photo of the real situation when a mouse was being trained.

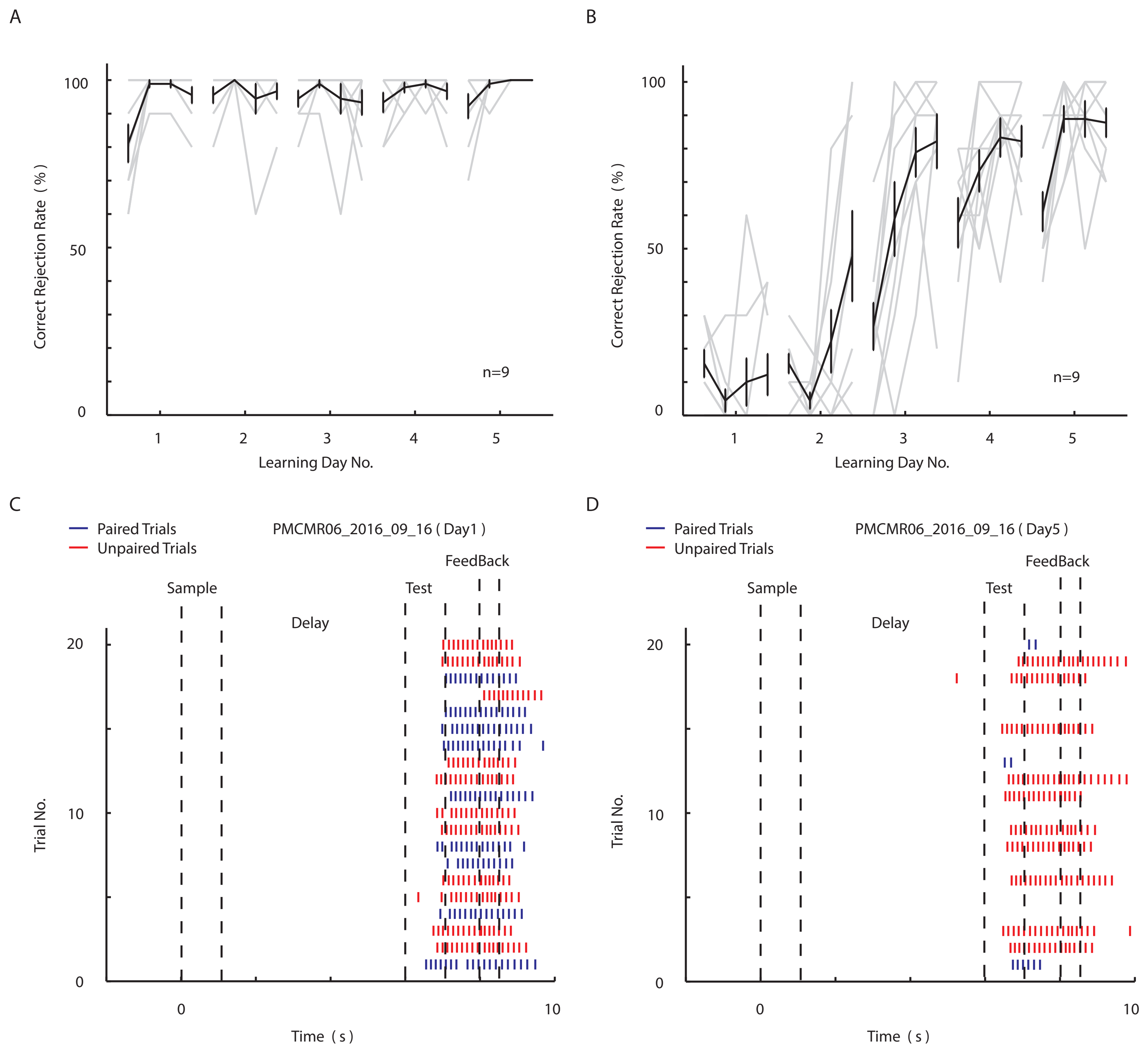


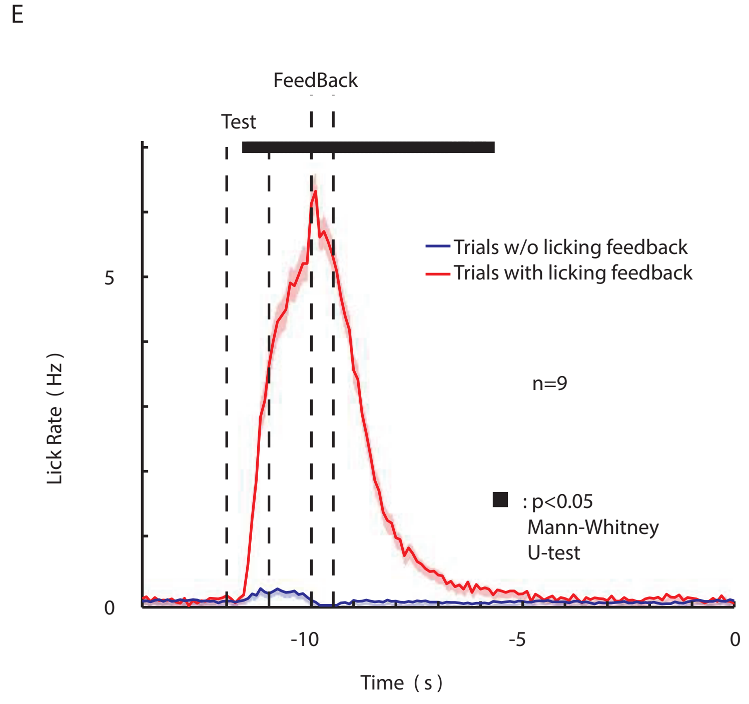


Supplementary Fig. 2. Details of the performance of mice during the learning of DPA task. (A) The hit rate curves throughout all 5 days of mice learning. (B) The correct rejection rate curves throughout all 5 days of mice learning. (C) The raster plot of licking in 20 trials of one mouse on the day1 of learning. (D) The raster plot of licking in 20 trials of the same mouse on the day5 of learning. (E) The histogram of licking rate from late delay to the onset of next trial sample period.

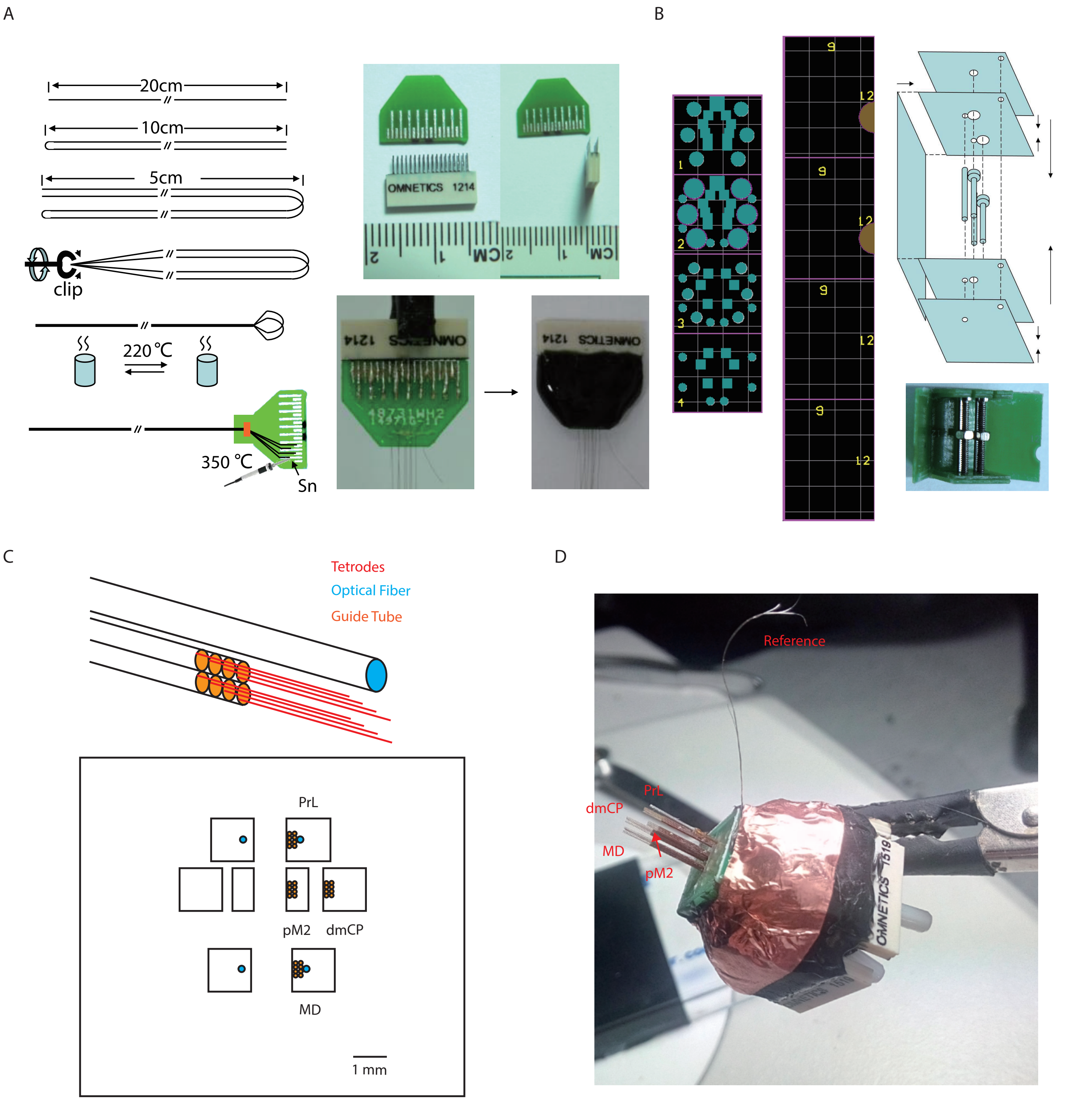


Supplementary Fig. 3. Protocol of making optrodes. (A) Procedure of making electrodes and connecting electrodes to interface. (B) Procedure of making micro-drive chamber.(C) Arrangement of electrodes and optical fibers. (D) Photo of a completed 128-channel multi-region recording optrode.

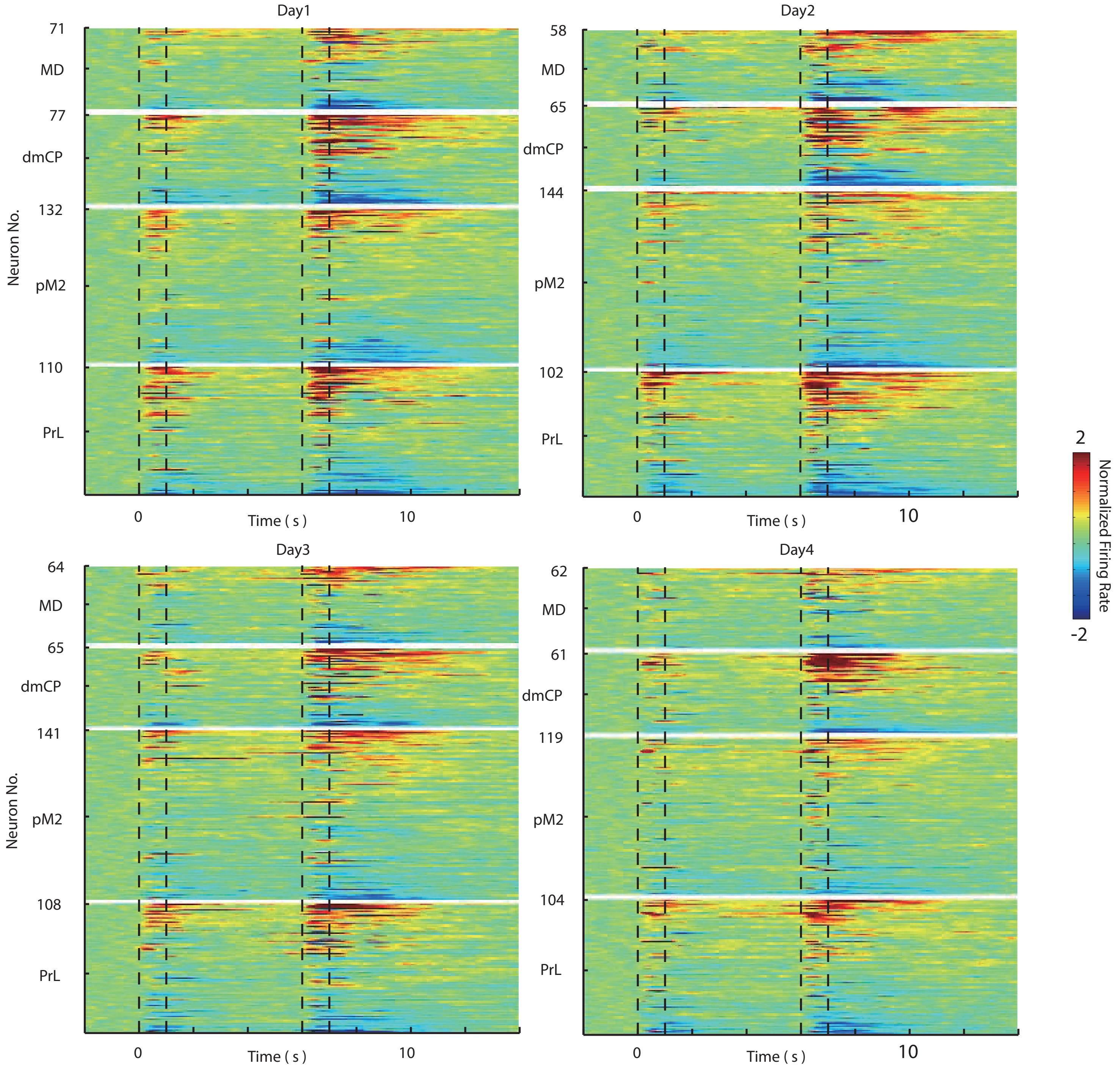


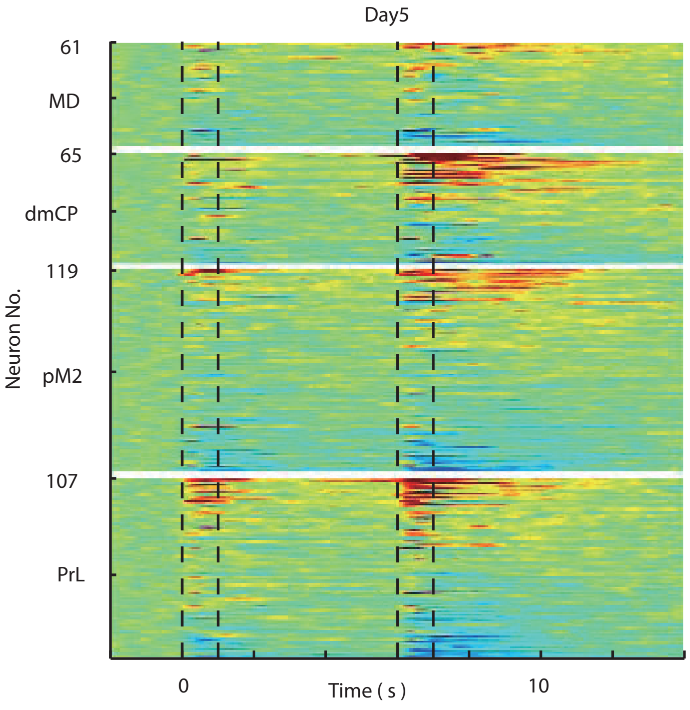


Supplementary Fig. 4. Heat maps for Neuronal activity of PrL, pM2, dmCP and MD on different learning days (day 1~5)

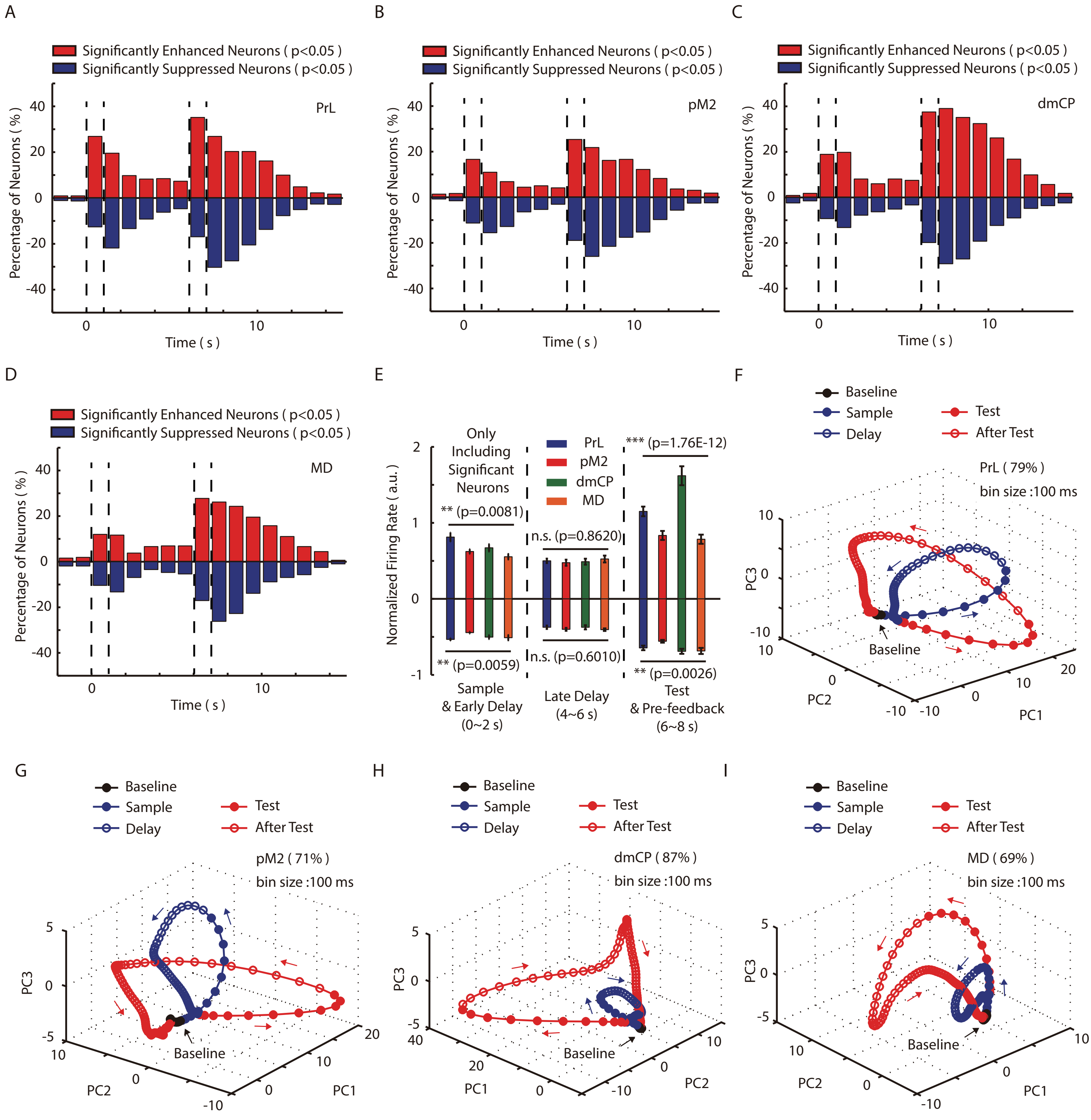


Supplementary Fig. 5. Details about the neural modulation in PrL, pM2, dmCP and MD during DPA task. (A) Percentage of PrL neurons significantly modulated according to baseline firing in each 100ms bin in DPA task. Red bars represent neurons enhanced. Blue bars represent neurons suppressed. (B) Similar as (A), but for pM2 neurons. (C) Similar as (A), but for dmCP neurons. (D) Similar as (A), but for MD neurons. (E) Comparing of neural modulation in PrL, pM2, dmCP and MD during sample period, late delay period and test period. Noted that the significant difference during sample and test periods. (F) PCA trajectories of neural dynamics in PrL during DPA task. (G) Similar as (F), but for pM2 neurons. (H) Similar as (F), but for dmCP neurons. (I) Similar as (F), but for MD neurons.

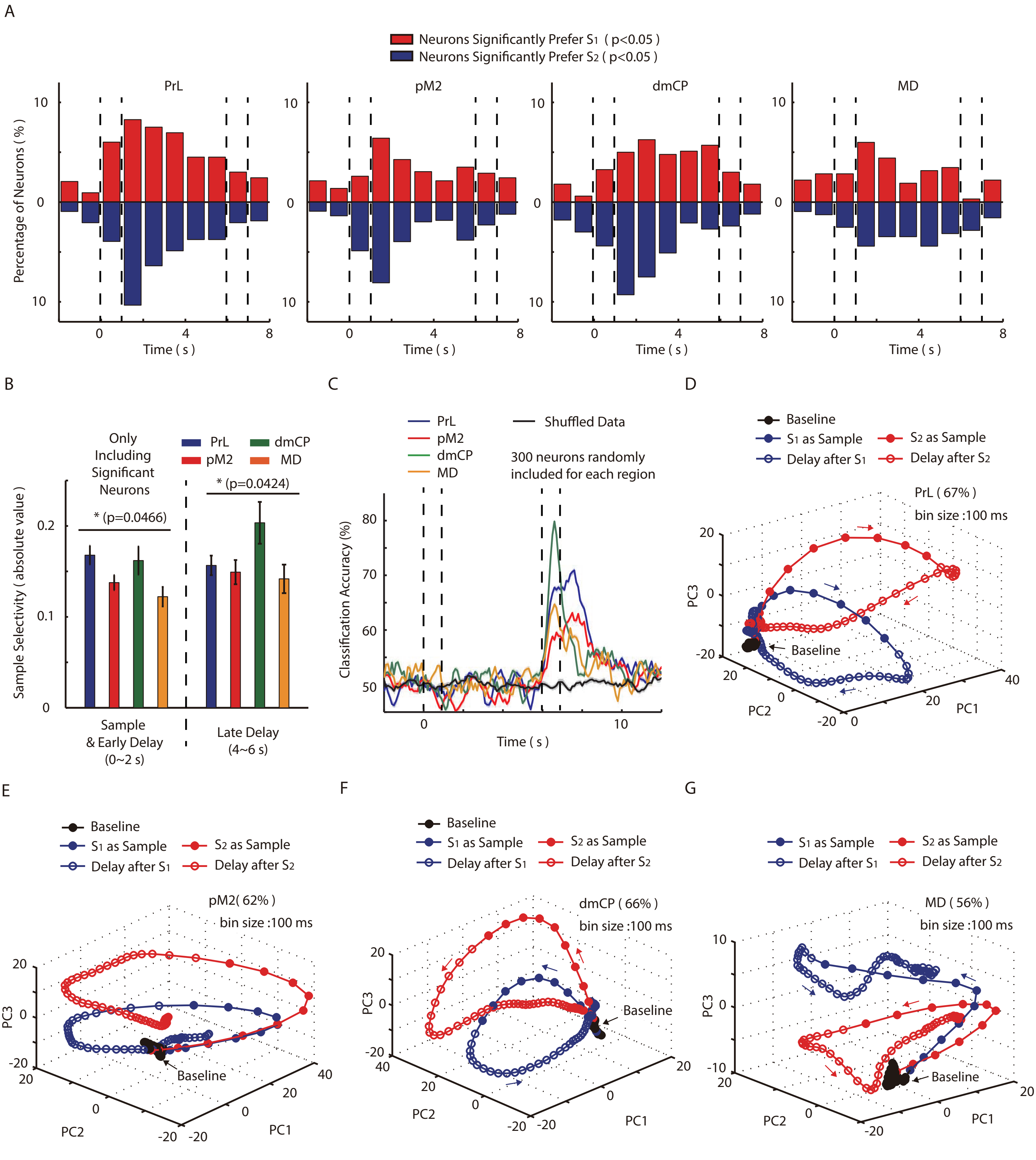


Supplementary Fig. 6. Details about the sample selectivity in PrL, pM2, dmCP and MD during DPA task. (A) Percentage of PrL, pM2, dmCP and MD neurons significantly selective to sample odors in each 100ms bin in DPA task. Red bars represent neurons preferring S1. Blue bars represent neurons prefering S2. (B) Comparing of sample selectivity in PrL, pM2, dmCP and MD during sample period and late delay period. (C) Decoding accuracy of neurons in PrL, pM2, dmCP and MD for test odors in DPA task. (D) PCA trajectories of neural dynamics in PrL in trails with S1 and trials with S2 during DPA task. (E) Similar as (D), but for pM2 neurons. (F) Similar as (D), but for dmCP neurons. (G) Similar as (D), but for MD neurons.

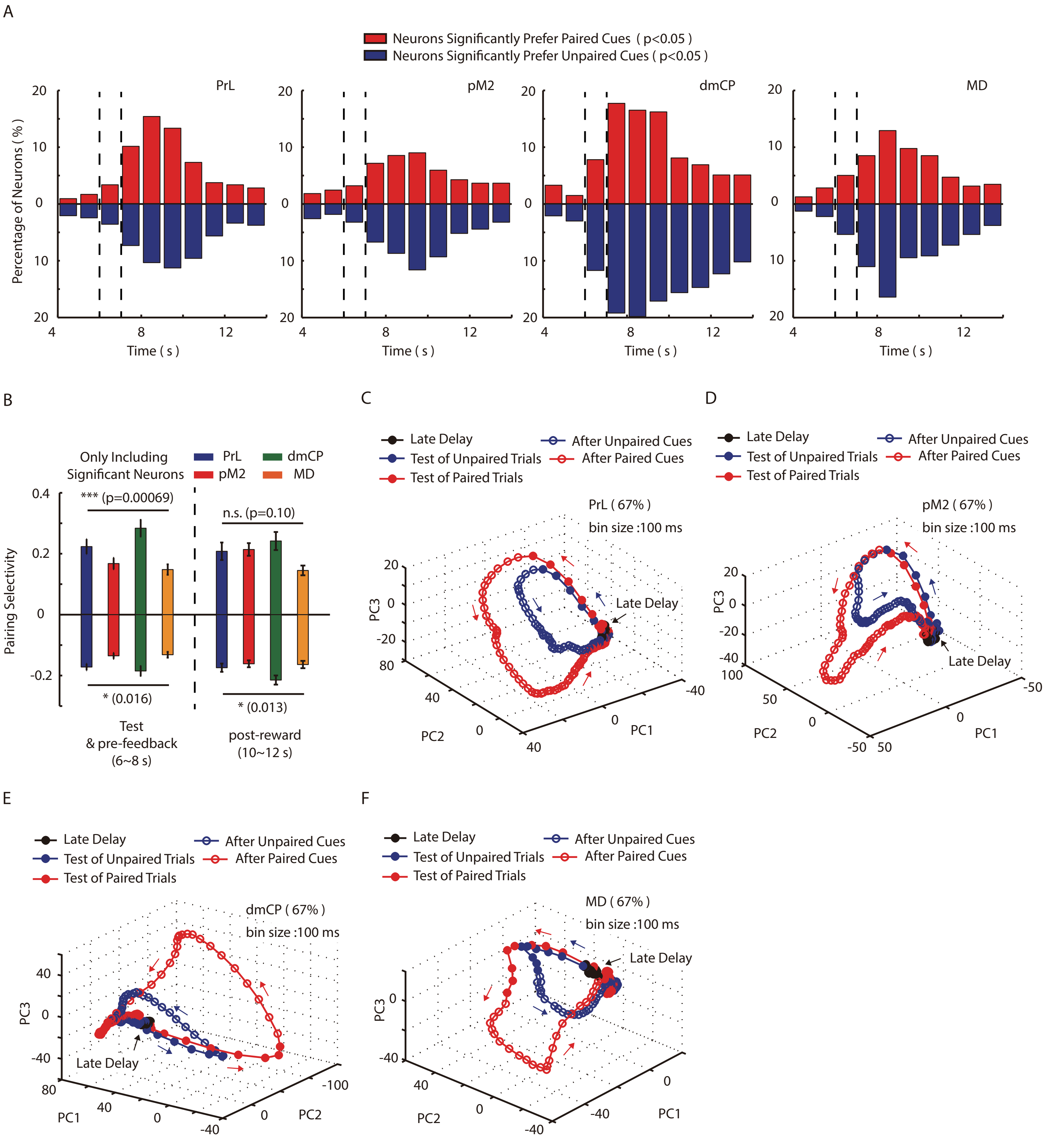


Supplementary Fig. 7. Details about the pairing selectivity in PrL, pM2, dmCP and MD during DPA task. (A) Percentage of PrL, pM2, dmCP and MD neurons significantly selective to sample-test pairing in each 100ms bin in DPA task. Red bars represent neurons prefering paired curs. Blue bars represent neurons prefering unpaired cues. (B) Comparing of pairing selectivity in PrL, pM2, dmCP and MD during test period and after reward feedback. (C) PCA trajectories of neural dynamics in PrL in trails with paired cues and trials with unpaired cues during DPA task. (D) Similar as (C), but for pM2 neurons. (E) Similar as (C), but for dmCP neurons. (F) Similar as (C), but for MD neurons.

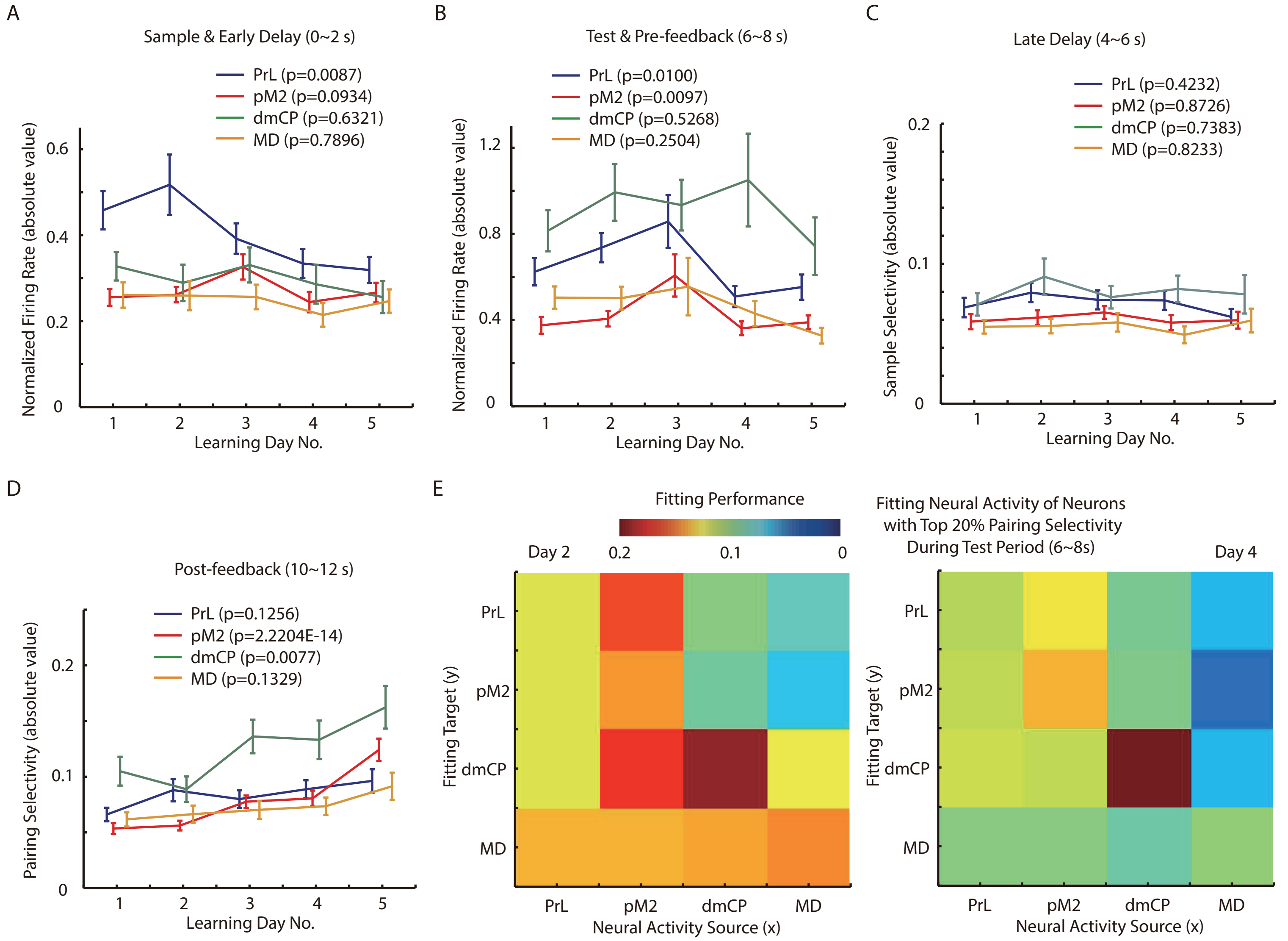


Supplementary Fig. 8. Details about the firing properties in PrL, pM2, dmCP and MD changing along DPA task. (A) Neural modulation during sample period in PrL, pM2, dmCP and MD in each day of learning DPA task. (B) Neural modulation during test period in PrL, pM2, dmCP and MD in each day of learning DPA task. (C) Sample selectivity during late delay period in PrL, pM2, dmCP and MD in each day of learning DPA task. (D) Pairing selectivity after reward feedback in PrL, pM2, dmCP and MD in each day of learning DPA task. (E) Fitting performance of neurons with top 20% pairing selectivity in PrL, pM2, dmCP and MD in day2 and day4 of learning DPA task.
